# Ubiquitous low-energy RNA fluctuations and energetic coupling measured by chemical probing

**DOI:** 10.64898/2026.01.28.702231

**Authors:** Edric K. Choi, Ritwika Bose, David H. Mathews, Anthony M. Mustoe, Julius B. Lucks

## Abstract

RNA function is governed by RNA folding, but strategies for measuring RNA folding thermodynamics are limited, and fundamental questions such as the energy of base pair opening remain debated. Here we introduce Probing-Resolved Inference of Molecular Energetics (PRIME) to extract nucleotide-resolution RNA structural energetics from scalable chemical probing experiments. Applying PRIME to diverse RNAs, we find that RNA base pairs and tertiary interactions dynamically open with free energies of 0.5-3 kcal/mol, revealing that RNA nucleotides ubiquitously sample open conformations at biologically accessible energies. PRIME further resolves energetic coupling across RNAs, providing an energetic understanding of RNA structural dynamics and long-range coordination in RNA folding. PRIME represents a widely accessible strategy for interrogating RNA thermodynamics, enabling mechanistic understanding and engineering of RNA biology.

## Main Text

RNA molecules have rugged thermodynamic folding free energy landscapes that critically impact every RNA biochemical process, including folding during transcription and splicing, translation by the ribosome, degradation, protein and ligand binding, and catalysis (*1-4*). Understanding RNA folding thermodynamics is thus central to understanding cellular biochemistry (*5*), the mechanisms of RNA-mediated diseases (*6*), and engineering RNA-based biotechnologies (*7*).

Despite decades of work, there continue to be major gaps in our understanding of the energetic principles governing RNA folding. Existing methods for measuring RNA thermodynamics suffer significant scale versus resolution tradeoffs. Even basic questions such as the energetic cost of disrupting a Watson-Crick-Franklin (WCF) base pair remain debated (*8*): measurements by tritium-exchange (H-T) (*9-12*) and UV melting experiments indicate Δ*G*^∘^≈ 1-3 kcal/mol (*13, 14*), whereas hydrogen-deuterium exchange (HDX) by NMR consistently report Δ*G*^∘^ ≈ 4-10 kcal/mol (*15, 16*). These disparate estimates translate into several orders of magnitude differences in how frequently an RNA base pair opens (3% for Δ*G*^∘^ = 2 kcal/mol, 0.0008% for Δ*G*^∘^ = 7 kcal/mol at 25°C), a fundamental structural transition involved in almost every RNA biological process (*4*). Even less is known about the energetics of non-canonical base-pairs, which are ubiquitous in functional RNAs and are critical to RNA three-dimensional folding (*17*). The inability to measure nucleotide-level energetics at scale poses a fundamental barrier to addressing these knowledge gaps and fully understanding and engineering RNA biology.

Here we present a new approach for measuring nucleotide resolution RNA folding thermodynamics through chemical <u>P</u>robing-<u>R</u>esolved <u>I</u>nference of <u>M</u>olecular <u>E</u>nergetics (PRIME). Chemical probing experiments use small molecule probes such as dimethyl sulfate (DMS) to preferentially modify unstructured RNA nucleotides, followed by measurement of the modifications by strategies such as mutational profiling sequencing (MaP) (*18*) (Fig. 1A). Probing experiments can be scaled to study tens of thousands of RNAs at a time and are widely used to measure RNA folding in cells (*19, 20*). Inspired by approaches to measure protein folding thermodynamics at residue-resolution with HDX (*11, 21*), PRIME uses a kinetic framework to extract quantitative, nucleotide-resolution energies 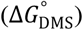 of RNA structural fluctuations from time-course DMS-MaP experiments. Application of PRIME to the *Salmonella* fourU RNA thermometer (*22*), the HIV-1 TAR hairpin (*23*), and the *Tetrahymena* P4-P6 domain (*24*), revealed that canonical base pairs dynamically fluctuate with energies of 1-3 kcal/mol, with non-canonical and tertiary pairs exhibiting comparable energies of 0.5-2.5 kcal/mol. These low energy RNA nucleotide fluctuations are expected to be populated significantly at physiological conditions and influence myriad RNA functions. We further show that PRIME uncovers coupling of nucleotide energetics across alternative RNA folds of the HIV-1 TAR hairpin, and compensation between energetics of secondary structure base pairs and tertiary interactions during Mg^2+^-dependent tertiary folding of the P4-P6 domain. PRIME bridges classical biophysics, chemical kinetics and high-throughput sequencing to make nucleotide-resolution RNA thermodynamics measurable at scale.

**Figure 1.**
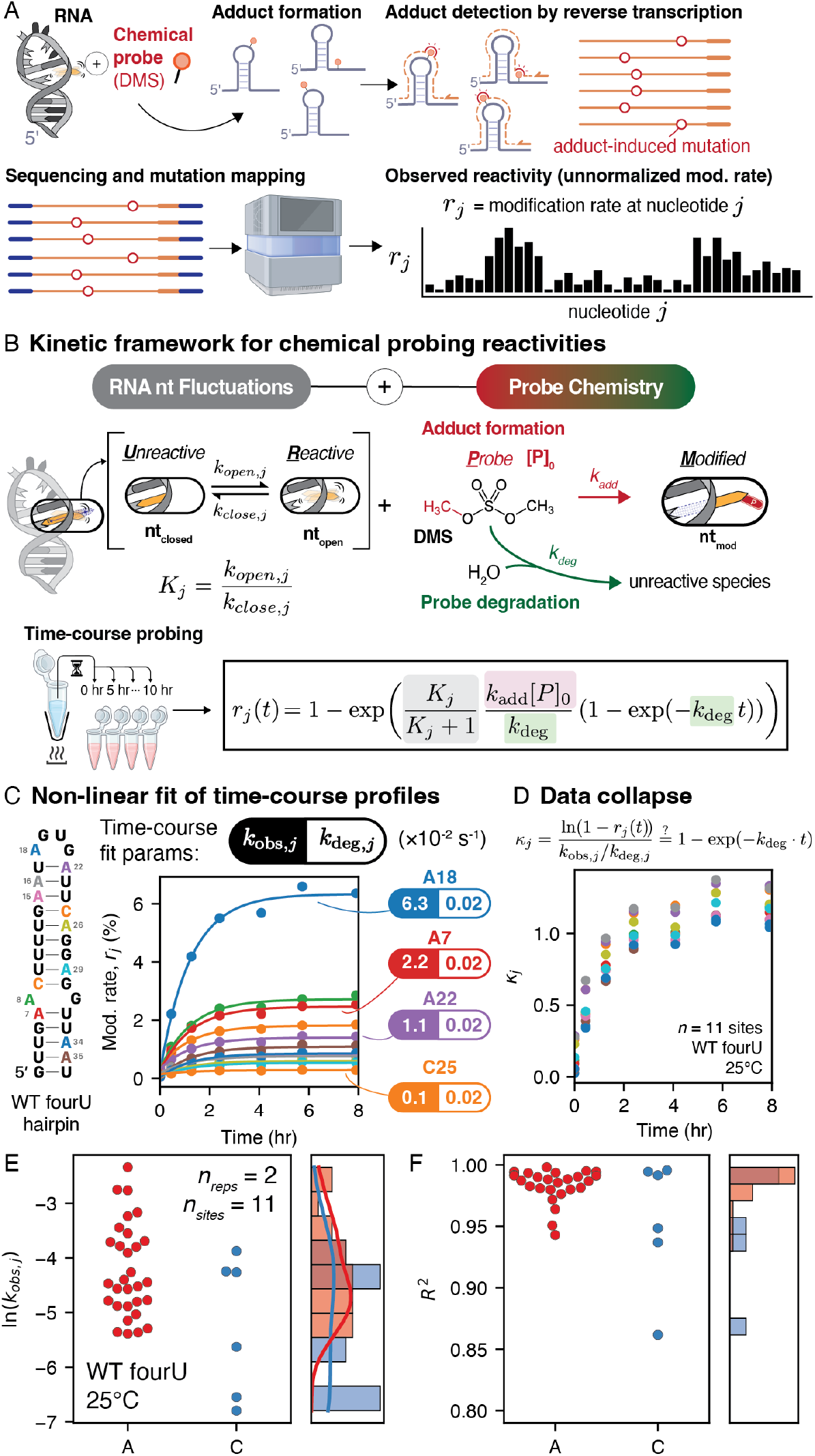
A kinetic model of RNA chemical probing accurately captures quantitative time-course DMS reactivities. (**A**) DMS probing experiments selectively modify RNA molecules at structurally accessible, reactive sites. Modifications are measured using mutational profiling by reverse-transcription and sequencing to yield a ‘reactivity’, *r*_*j*_, for each nucleotide *j*, representing the modification fraction at that position. (**B**) RNA nucleotides fluctuate between reactive and non-reactive configurations, characterized by an equilibrium constant of fluctuations, *K*_*j*_. A quantitative kinetic model shows that reactivity depends on intrinsic information about RNA structure (*K*_*j*_), and experimental parameters ([*P*]_0_, *k*_add_, *k*_deg_, *t*). (**C**) Time-course DMS probing of the WT fourU RNA thermometer at 25°C yields kinetic reactivity traces that are accurately fit by the model using two constants, *k*_obs,*j*_ and *k*_deg,*j*_. Representative data for all adenosine (A) and cytidine (C) sites are shown. (**D**) Reactivity time courses collapse onto a common curve when transformed to isolate *k*_deg_ according to Eq. (1). (**E-F**) Distribution of ln(*k*_obs,*j*_) and R^2^ values from non-linear regression of time-course probing data (three replicates, 25°C) for A and C nucleotides. Histograms share a common y-axis, with corresponding density curves shown on the right.

### Time-dependent DMS probing obeys a quantitative HDX-like chemical kinetics model

We built on HDX theory (*21*) to develop a quantitative model of the RNA chemical probing reaction (Fig. 1B). In this model, each RNA nucleotide, *j*, fluctuates between an unreactive state (*U*_*j*_) and reactive state (*R*_*j*_), with rate constants *k*_open,*j*_ and *k*_close,*j*_. The chemical probe *P* reacts irreversibly with *R*_*j*_ to form a modified state *M*_*j*_ at an adduct formation rate *k*_add_, while *P* concurrently undergoes hydrolytic degradation at rate *k*_deg_ (Fig. 1B). Under typical conditions, local RNA fluctuations are much faster than probe chemistry such that *k*_open,*j*_, *k*_close,*j*_ ≫ *k*_*add*_[*P*]_0_, *k*_deg_ (*25-27*), rendering each site pre-equilibrated prior to reaction, corresponding to an EX2 (bimolecular) regime in HDX. This regime yields an analytical expression for observed reactivity as a function of reaction time, *r*_*j*_(*t*):

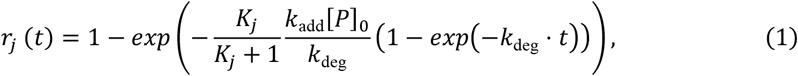

where *K*_*j*_ = *k*_open,*j*_/*k*_close,*j*_ is the fluctuation equilibrium constant (Supplementary Note 1). Exact numerical simulations confirmed that Eq. (1) accurately captures nucleotide modification kinetics over a broad range of opening and closing rates (fig. S1), breaking down only under unrealistically high RNA concentrations (fig. S1B) or when structural transitions are very slow (>100 s) (fig. S1C). Eq. (1) makes clear that reactivity is a nonlinear convolution of intrinsic RNA structure parameters (the equilibrium constant *K*_*j*_ of fluctuations between open and closed states) and variable experimental parameters ([*P*]_0_, *k*_add_, *k*_deg_, *t*), limiting interpretability of typical single time-point reactivity measurements. Importantly, Eq. (1) provides a framework for directly extracting *K*_*j*_ from probing kinetics, which forms the basis of the PRIME approach.

To validate the predicted kinetic behavior of DMS modification, we performed time-course DMS probing of the fourU RNA thermometer hairpin, a well-characterized model RNA that regulates translation of the *agsA* gene in *Salmonella* (*22, 28*) (Fig. 1C). Probing was performed at 25°C on *in vitro* transcribed fourU RNA using buffers optimized to maintain constant pH over the duration of the reaction (fig. S2) (*29*), and probe concentrations optimized to yield single-hit reaction conditions (fig. S3). Time resolution was achieved by quenching aliquots of the reaction across six or more time points over 8 hours (Methods). The modification fraction at each nucleotide and each time point, *r*_*j*_(*t*), was then measured by MaP reverse-transcription and sequencing (*30*). The resulting reactivity time-courses for adenosines (A’s) and cytidines (C’s), which are most robustly modified by DMS, demonstrated rapid initial increases that gradually plateaued as the probe gets depleted, as predicted by Eq. (1) (Fig. 1C). Individual nucleotide reactivity traces were fit to Eq. (1) to extract two parameters: *k*_obs,*j*_ = [*K*_*j*_/(*K*_*j*_ + 1)*k*_add_[*P*]_0_, and *k*_deg_ (Fig. 1C). Consistent with the expected secondary structure, the unpaired nucleotide A18 exhibited large *k*_obs,*j*_ values (0.06 s^-1^), corresponding to a higher plateau point, while base-paired nucleotides show 2 orders of magnitude lower *k*_obs,*j*_ and reached lower plateau values (Fig. 1C). As predicted by Eq. (1), transformation of the data revealed a collapse of all time course reactivity traces onto a single curve (Fig. 1D), consistent with a shared value of *k*_deg_ across nucleotide positions as expected from the probe chemistry. Accordingly, *k*_deg_ was fit globally across sites with no measurable loss in fit quality (fig. S4). Across several orders of magnitude in *k*_obs,*j*_ (Fig. 1E), we found fits to Eq. (1) were robust, with all of nucleotides in WT fourU achieving R^2^ > 0.85 at 25°C (Fig. 1E-F) and showing high reproducibility across independent replicates (fig. S5). These results demonstrate that chemical probing data are quantitatively governed by Eq. (1) and yield internally consistent rate parameters.

### DMS reaction rates from time-course probing match those from NMR

We next sought to validate that kinetic rates measured by PRIME are physically meaningful. To test expected physical relationships between temperature, folding, and modification kinetics, we repeated time-course DMS probing of the fourU RNA across a broad temperature range (Fig. 2A). Consistent with our analysis at 25°C, these additional time-course data demonstrated excellent fits to Eq. (1) (fig. S6).

**Figure 2.**
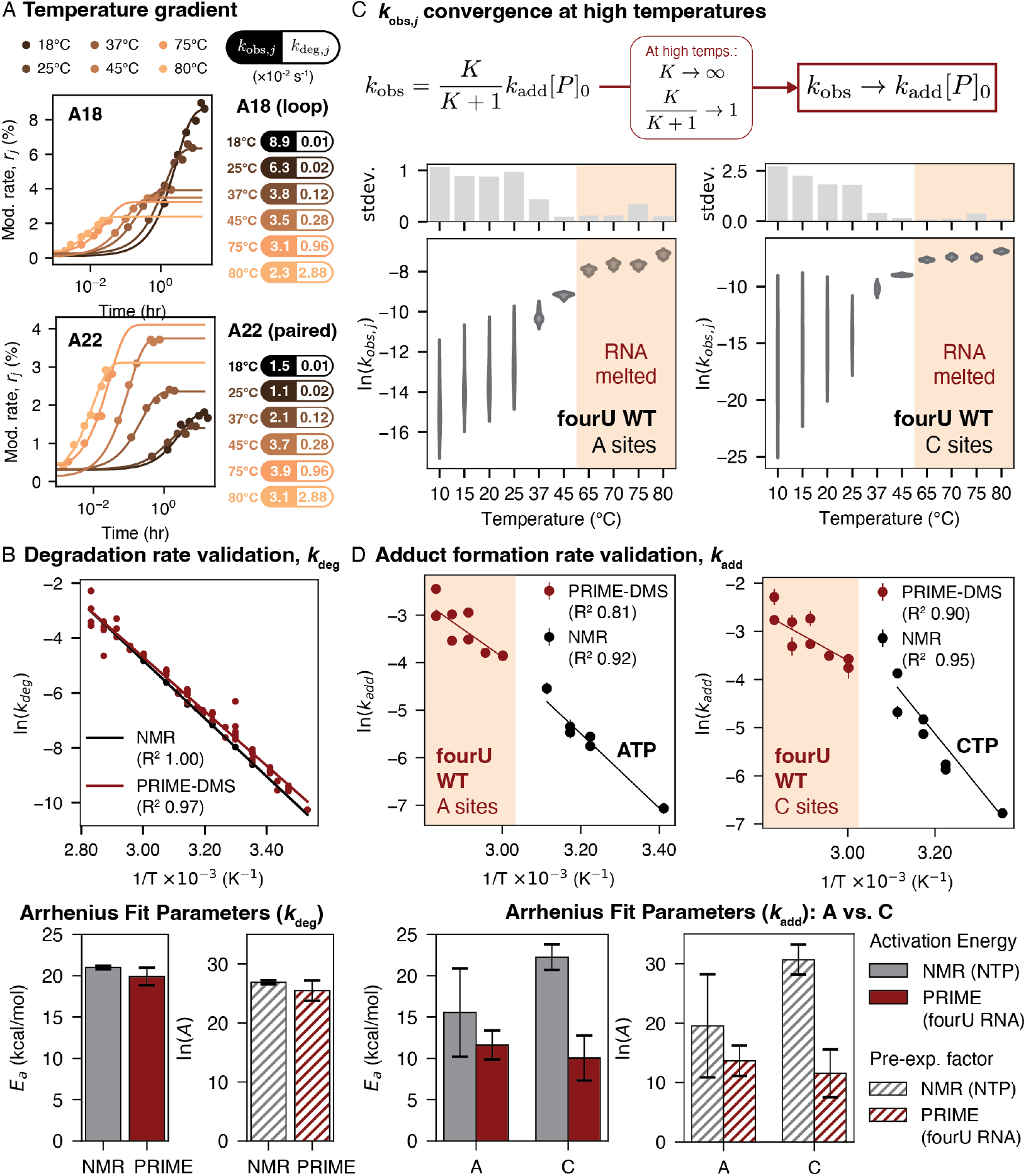
NMR validation of PRIME-derived degradation and adduct formation rates. (**A**) Representative time-course plots for nucleotides A18 and A22 in WT fourU across select temperatures; each time-course yields *k*_obs,*j*_ and *k*_deg_ quantities, which are listed on the right. (**B**) Arrhenius analysis of ln *k*_deg_ comparing independent NMR measurements with globally fitted PRIME-DMS rates from the fourU RNA. Bar plots summarize Arrhenius fit parameters (Methods). (**C**) *k*_obs_ collapses to the adduct-formation term *k*_add_[*P*]_0_ at high temperatures. Violin plots for 8 adenosine sites and 2 cytidine sites show convergence of *k*_obs,*j*_ values toward a common limiting rate at high temperature. Red shading indicates temperatures at which the RNA is expected to be unfolded based on the global melting temperature (*28*). Accompanying bar plot shows the standard deviation of ln *k*_obs,*j*_ across sites at each temperature. (**D**) Arrhenius analysis of ln *k*_add_ values grouped by base (A/C) extracted from panel (C) (red) compared to independent NMR measurements (black, fig. S8 for details). Bar plots summarize Arrhenius fit parameters (Methods), error bars represent standard error from linear regression.

*k*_deg_ rates measured by PRIME demonstrated logarithmic dependence on 1/T (Fig. 2B), confirming expected Arrhenius behavior of DMS degradation with water. Significantly, these *k*_deg_ rates were in quantitative agreement with independent ^1^H NMR measurements of DMS degradation in matched buffer conditions (Fig. 2B, fig. S7).

*k*_obs,*j*_ represents a convolution of the reactive state equilibrium constant, *K*_*j*_, and the adduct formation rate, *k*_add_[*P*]_0_. At low temperatures, base paired nucleotides should have small *K*_*j*_ and thus low *k*_obs,*j*_, resulting in disperse *k*_obs,*j*_ between paired and unpaired bases. Indeed, *k*_obs,*j*_ sharply decreases and exhibits progressively greater dispersion at lower temperatures (Fig. 2C). In contrast, as the RNA melts, *K*_*j*_ → ∞ and *k*_obs,*j*_ should converge toward a common value across nucleotides, corresponding to *k*_add_[*P*]_0_. Indeed, *k*_obs,*j*_ collapsed to a uniform rate at high temperatures, consistent with the modification reaction becoming chemically rate limited (Fig. 2C, bottom; fig. S9). Furthermore, these melted *k*_obs,*j*_ values follow an Arrhenius temperature dependence, as expected for *k*_add_ (Fig. 2D). To independently validate these PRIME *k*_add_ measurements, we used time-resolved ^1^H NMR experiments to measure *k*_add_ from DMS methylation of free ATP and CTP nucleotides (fig. S8). Due to time-resolution constraints, NMR *k*_add_ data was only obtainable at low temperatures. Nonetheless, Arrhenius analysis of NMR data yielded activation energies and pre-exponential factors within error of PRIME values for A nucleotides. Measurements for C’s were comparable magnitude but were approximately 2-fold higher by NMR, which may reflect increased uncertainty in the PRIME-derived estimates due to the small number of C residues (*n* = 2) in the fourU construct or modest chemical differences in reaction with free CTP versus polynucleotides.

Together, these results demonstrate that PRIME recovers physically meaningful kinetic rates of chemical probing, physically grounding our kinetic framework, and establishing an approach to deconvolve nucleotide-level RNA energetics from chemical probing data.

### PRIME-DMS reveals a low energetic cost of RNA base pair opening

Having validated Eq. (1), we next used it to extract the free energy of the unreactive/reactive structural equilibrium that governs the reactivity of each nucleotide. Specifically, we used PRIME-fit *k*_deg_ rates, PRIME-fit *k*_add_ rates and known [*P*]_0_ to compute the per-nucleotide equilibrium constant, *K*_*j*_ (Supplementary Note 2; Methods). Here, *K*_*j*_ reflects the equilibrium of a nucleotide between a structurally constrained unreactive state such as a base pair, and a reactive state accessible to DMS attack such as opening of the pair. *K*_*j*_ can also report on other structural constraints such as nucleotide stacking. We then calculated per-nucleotide DMS probing free energies, 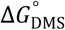 as (Supplementary Note 3-4):

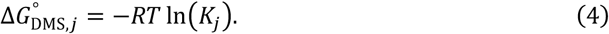

At 25°C, base-paired positions in the fourU hairpin exhibit free energies in the range 1-4 kcal/mol with C-G sites in the range 3.4-4.2 kcal/mol and A-U sites in the range 0.95-2.0 kcal/mol (Fig. 3A). These values align with canonical estimates of ∼0.5-1.5 kcal/mol per hydrogen bond inferred from UV melting of mutated short hairpin variants (*31*). The WT RNA features a A8-G31 pair (1.2 kcal/mol) which is comparable in stability to the neighboring A7-U32 but weaker than the loop-closing A22-U17 pair (1.7 kcal/mol). In contrast, unpaired A18 exhibits 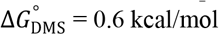, consistent with nucleotide stacking. These 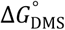 measurements were highly reproducible across independent replicates (fig. S10).

**Figure 3.**
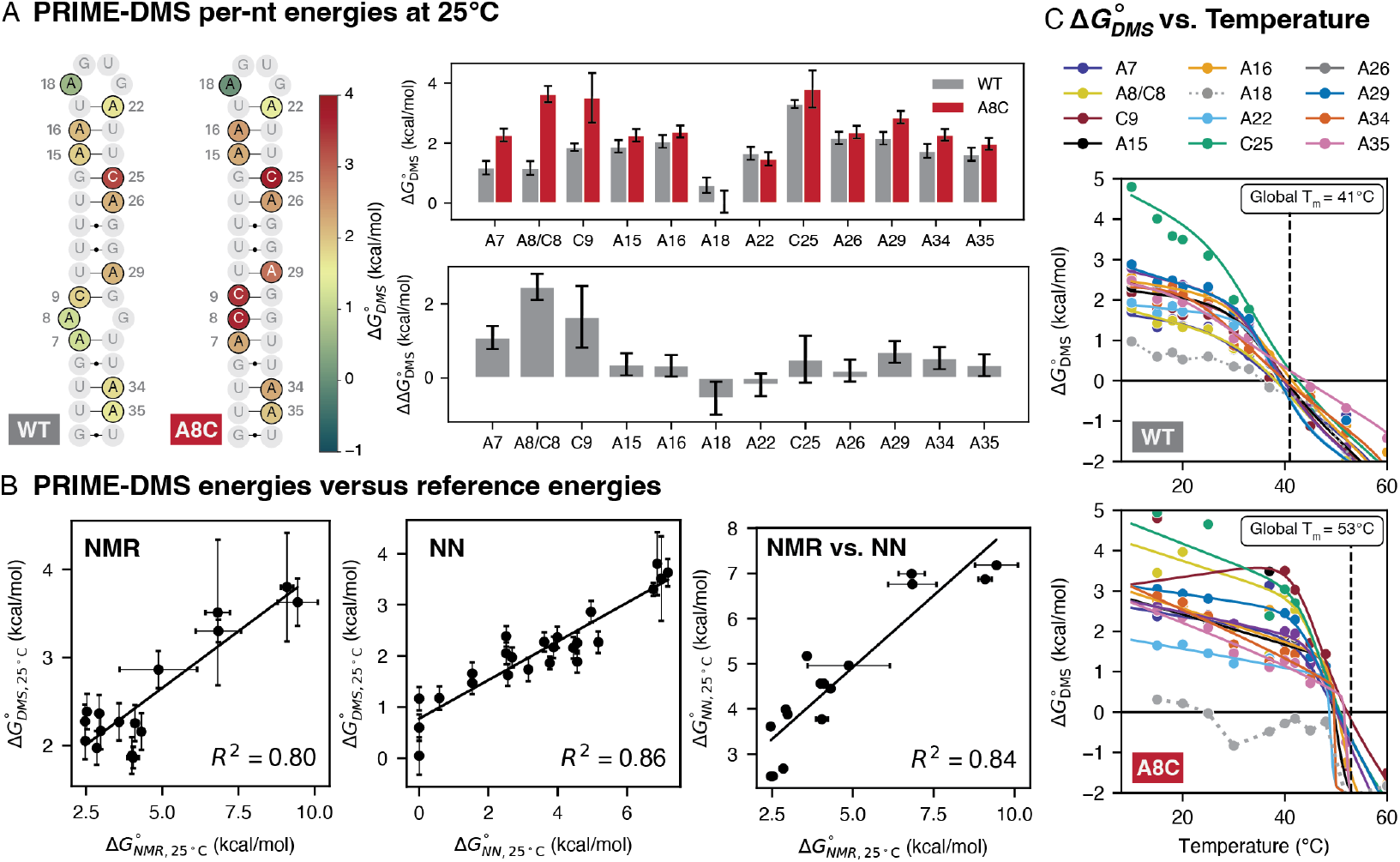
PRIME reveals low energy per nucleotide RNA structural fluctuations. **A**, left) Secondary structure of WT and A8C fourU thermometer RNAs colored by 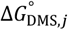 values for A and C positions. Nucleotides without color were excluded from analysis due to low-confidence fits. (**A**, right) PRIME-derived free energies (top) and differences between WT and A8C variants (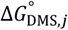, bottom) at 25°C. (**B**) Linear regressions comparing 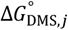 against reference base pair dissociation energies from NMR imino proton exchange, nearest-neighbor (NN)-based ensemble calculations, and NMR/NN correlation. PRIME experiments are performed at matching buffer conditions as NMR experiments. (**C**) Temperature-dependence plots of 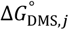 quantities for WT (top) and A8C (bottom) fourU thermometer. Solid lines are transformed from 2-state melting curve fits of ln(*k*_obs_) (fig. S9). For A18, a dotted line connects data points because the data could not be fit to a two-state model. Global melting temperature obtained from CD melt experiments performed in (*28*) are indicated as a black dashed vertical line. 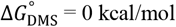 is indicated as a solid horizontal aixs. All error bars are propagated from standard error from kinetic fits and 3 biological replicates.

We also studied the A8C mutant, which replaces the central A-G mismatch with a canonical C-G base pair (*28*). This mutation increases the stability of immediately adjacent base pairs by ∼1-2 kcal/mol (Fig. 3A), consistent with enhanced local stacking and improved helical continuity This increased stability further propagates several base pairs both up and down the stem 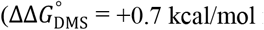 for A29 and +0.5 kcal/mol for A34). Interestingly, A18 becomes modestly destabilized in the A8C mutant 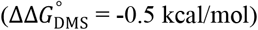, suggesting that local helix stabilization can propagate energetic effects to distal loop regions.

We focused on the fourU RNA because its thermodynamics have also been characterized by NMR imino-proton exchange experiments (*28*). Direct comparison between 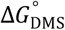 values for A and C positions and corresponding NMR-derived free energies for U and G positions in the same base pairs revealed remarkably strong correlation across both WT and A8C mutants (R^2^ = 0.80; Fig. 3B). 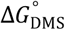 is similarly strongly correlated with energies predicted from the nearest-neighbor (NN) energy model (*32*), 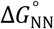, which we calculated as the energetic penalty of forced base pair disruption (R^2^ = 0.86; Fig. 3B; Methods). These PRIME correlations are equivalent to the correlation between NN and NMR energies (R^2^ = 0.84; Fig. 3B), indicating that PRIME-derived energetics capture the shared physical basis of these models while resolving features unique to each. By contrast, conventional endpoint DMS reactivities were uncorrelated with NMR energies, either as raw reactivities or transformed into pseudo-free energies, as conventionally used for RNA structure modeling (R^2^ = 0.01; fig. S12).

While being strongly correlated, 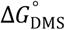 values are ∼50% lower than NMR-and NN-derived free energies (1-4 kcal/mol versus 2-10 kcal/mol; Fig. 3B). Given that our DMS buffers were matched to NMR buffers, whereas NN energies derive from very different buffer conditions, this discrepancy is not explainable by environmental variables. Rather, this discrepancy likely reflects differences in the structural states measured by each method. NMR-detected imino proton exchange is a two-step reaction that requires interactions with an external base catalyst, which may require more extensive base-pair disruption to reach the reactive state. Similarly, NN energies report on complete base-pair melting rather than more subtle base pair fluctuations. In contrast, our data suggest that DMS methylates lower-energy base-pair-opened states that likely retain partial stacking or hydrogen bonding. Notably, our 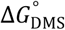 values closely match classic tritium-exchange measurements on DNA helices (∼2–3 kcal/mol) (*9, 10, 12*). Thus, PRIME supports that RNA base-pair opening only incurs a modest energetic penalty.

The ability to quantify local 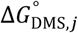 at nucleotide resolution also enables direct connection between site-specific energetics and global melting behavior. At low temperatures, when the population of the unfolded state is low, each nucleotide exhibits distinct 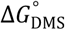, reflecting individual opening events within the folded helix (Fig. 3C). However, as the RNA nears the global melting temperature, 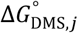 of paired residues collapse to a single melting curve, indicating transition to a regime where 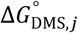 reports on global unfolding. Indeed, the nucleotides converge to 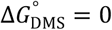 at precisely the independently measured melting temperature (*28*) for the WT (41°C), and within uncertainty for A8C variants (53°C) owing to noisier fits (Fig. 3C). A steeper approach to Δ*G* = 0 in the A8C variant is consistent with enhanced folding cooperativity due to its longer uninterrupted stem (Fig. 3C). As a notable exception, the unpaired nucleotide A18 follows a distinct “melting” trajectory and crosses 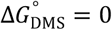 at a lower temperature than paired residues. A18 further demonstrates a lower ‘melting’ temperature in A8C than in WT (∼20°C vs ∼30°C) despite having an identical local sequence context, again consistent with apical loop energetics varying due to altered helical structure in the A8C variant.

Overall, these data demonstrate that nucleotide-resolved 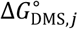 values report on a distinct and physically meaningful regime of RNA base-pair opening that is not captured by conventional endpoint reactivities, nearest-neighbor models, or NMR imino-proton exchange alone. These DMS-accessible opened states likely retain partial stacking interactions and are significantly populated well below melting temperature, placing them on the same energetic scale as classic tritium-exchange measurements. The convergence of 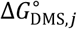 to zero at the global melting temperature further establishes that local opening energetics are directly coupled to cooperative RNA unfolding. More broadly, these findings show that chemical probing kinetics can be leveraged to extract quantitative, nucleotide-level energetic information that bridges local dynamics and global folding thermodynamics.

### Secondary structure population dynamics induce long-range nucleotide energetic coupling

Many RNAs dynamically transition between alternative base-pairing arrangements (*4*), but the energetics governing these dynamics remain incompletely understood. To investigate the relationship between nucleotide-level energetics and global structural dynamics, we applied PRIME to the well-studied TAR stem-loop of HIV-1 (*33-38*) (Fig. 4A). TAR has been shown to populate at least three alternative paired states: a dominant ground state (GS, 87%), an excited ES1 (12%) state featuring alternative pairing within the TAR apical loop, and ES2 (<1%), which features a base-pairing register shift and formation of multiple non-canonical pairs in the upper SL2 stem (Fig. 4A). While global TAR dynamics have been extensively studied by NMR, nucleotide-level energetics of TAR have not been resolved.

**Figure 4.**
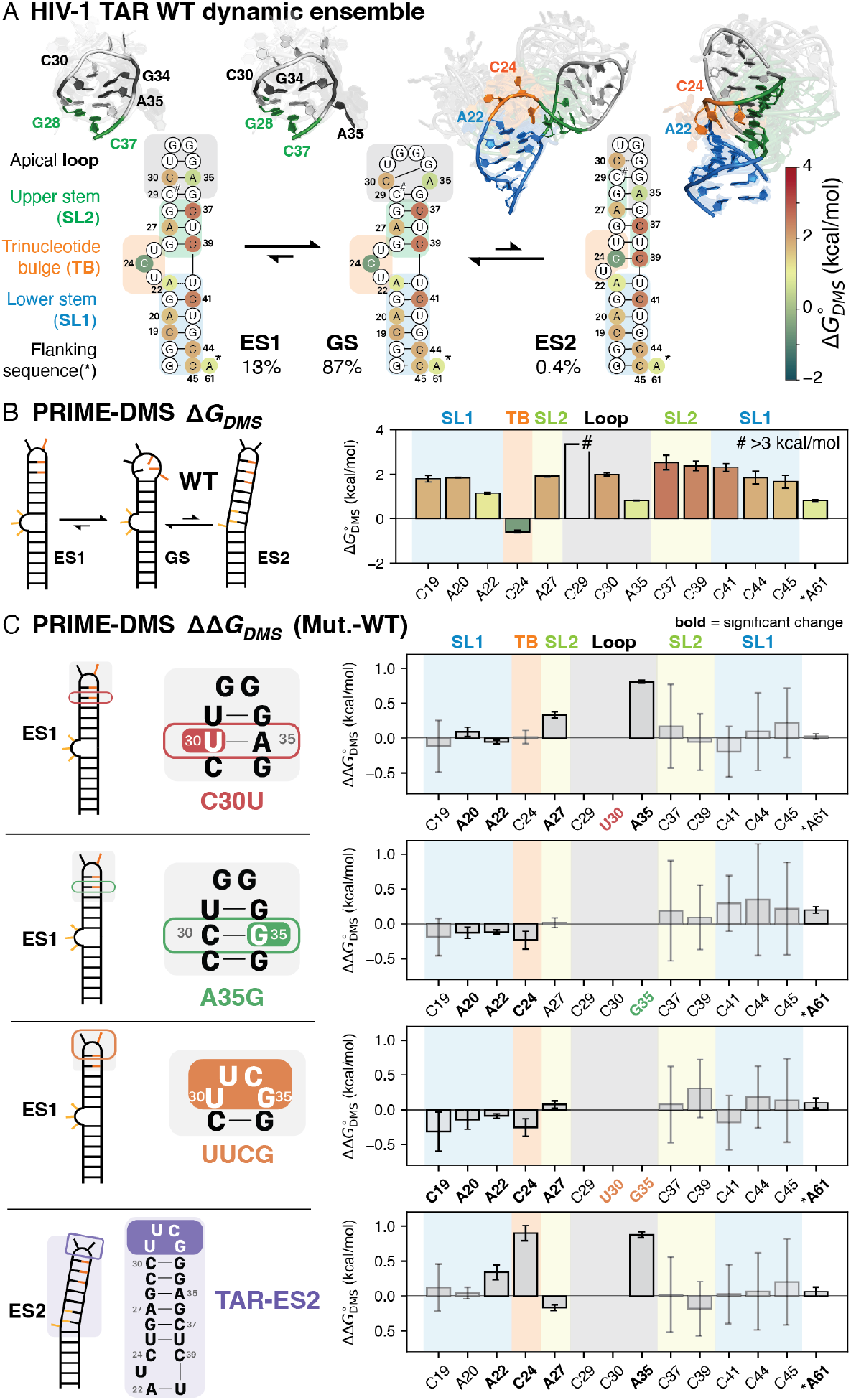
PRIME-DMS nucleotide energetics reveal population-mediated energetic coupling in HIV-1 TAR. (**A**) The WT HIV-1 TAR hairpin samples a ground state (GS) and two excited states (ES1, ES2). NMR-derived secondary structures for the different states are color-coded according to 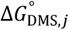, alongside representative GS, ES1, and ES2 NMR-derived ensembles (*39, 40*); apical loop ensembles for ES1 and GS are color coded according to nucleotide positions, hairpin ensembles for GS and ES2 are color coded according to annotated regions. (**B**) PRIME-derived free energy 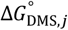 bar plot for WT TAR construct. Bar highlighting is according to the annotated regions in (A). The bar marked with a hash symbol (#) denotes a site with reactivity too low to be reliably fit by the time-course model and can be interpreted as having 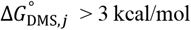 (fig S13). A61 is marked with a asterisk (*) correspond to unpaired flanking sequences not shown in structures. Data represent averages over two independent time-course replicates; error bars denote propagated standard error from time-course fits across replicates. (**C**) 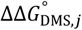 of TAR mutants relative to WT (Mut.-WT) that bias the conformational ensemble toward specific structural states (in parenthesis): C30U (ES1), A35G (ES1), UUCG (ES1), TAR-ES2 (ES2). 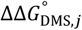 values were considered significant when error bars excluded zero; sites meeting these criteria are highlighted in **bold**. Note that the UUCG and TAR-ES2 constructs introduce a two-nucleotide deletion and insertion in the loop, respectively, but residue numbering was adjusted to match the WT sequence. Secondary structures are presented alongside per-nt 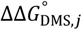 profiles.

We collected time-course PRIME-DMS data on WT TAR at 25°C (Fig. 4B), with data fitting reproducibly to the PRIME framework across two independent replicates (fig S10). We obtained high quality fits for all A positions and 8/9 C positions (fig. S13A). Base pairs in SL1 and SL2 exhibited 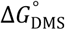 values of ∼1.5-3 kcal/mol, comparable to the base pair energies observed in the fourU system. The nominally paired residue A22 exhibited lower 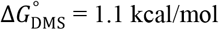, consistent with local fraying observed by NMR (*37*) (Fig. 4A). In the apical loop, C30 exhibited 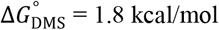, consistent with expected cross-loop pairing in the GS and nearing the stability of a standard helical pair. Unpaired nucleotide A61 exhibited 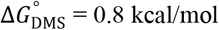, consistent with DMS accessibility being modulated by base-stacking. By contrast, C24 in the UCU bulge displayed a negative 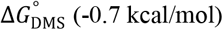, indicating it is preferentially in a reactive conformation, consistent with NMR observations that C24 is oriented away from the helix in most three-dimensional conformations of the GS (Fig. 4A) (*39, 40*).

As WT TAR RNA exists in an ensemble of states, analysis of WT energetics alone is unable to reveal contributions from individual states. To address this limitation, we measured 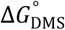 for TAR mutants designed to stabilize the ES1 and ES2 states and then computed the per-nucleotide change in stability relative to WT (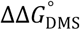; Fig. 4C, fig. S14) (*35, 37, 38*). While many positions showed minimal 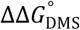, several positions showed significant changes indicative of energetic redistribution within each mutant. In the C30U mutant, which is expected to stabilize ES1 without impacting ES2, changes were local: A35 exhibits 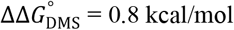, consistent with compaction of the apical loop; and A27 exhibits 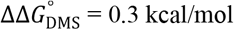, indicating propagated stabilization of SL2. By contrast, the A35G and UUCG mutants destabilize ES2, and both exhibit distal changes in stability at A20, A22 and C24 (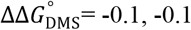, and -0.2 kcal/mol, respectively). This destabilization is consistent with these mutants favoring the open bulge structure of ES1. Indeed, the ES2-stabilizing mutant TAR-ES2 showed a dramatic stabilization of position C24 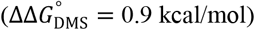 and A22 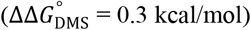 (Fig. 4C), consistent with non-canonical C24-C39 pairing and stabilization of A22 by SL1-SL2 interhelical stacking (Fig. 4A). TAR-ES2 also features modest destabilization of A27 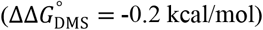, consistent with replacement of a WCF pair in WT/ES1 by a GA pair. Notably, A35 also forms a GA pair yet displays a substantially larger 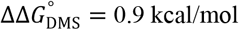, underscoring strong context-dependence of non-canonical interactions. Thus, ES2 provides significant net-stabilization of nucleotides throughout TAR, helping rationalize why the non-canonical ES2 secondary structure is populated by the WT sequence.

Together, these results reveal that secondary structure dynamics can couple nucleotides energetics across comparatively long distances, and correspondingly that nucleotide energetics encode information about population shifts within multi-state RNAs.

### PRIME resolves secondary-tertiary energetic coupling at nucleotide resolution

Much of RNA structural and functional diversity derives from the ability of RNAs to fold into complex 3D structures (*4, 41*). However, the fluctuations that occur within these 3D structures and the relationship between nucleotide-level energetics and global 3D folding remains largely unknown. Large RNAs are challenging to study by NMR (*42*), while existing biophysical methods report on global thermodynamics and lack nucleotide resolution (*43-45*).

We applied PRIME to obtain the first nucleotide-resolution energetic map of a large tertiary folded RNA, using the canonical model P4-P6 domain of the *Tetrahymena* group I intron ribozyme (*24*). Time-course DMS experiments were performed at 23°C and 5 mM MgCl_2_, matching established experimental conditions (*46*). Modification kinetics were well-fit by PRIME, yielding reproducible per-nucleotide 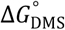 values for all 49 adenosine and 21/35 cytidine sites (Fig. 5A; fig. S13B, S15). The 14 unmeasurable cytidines were all base-paired and displayed reactivities too low for reliable kinetic fitting, yielding lower-bound estimates of 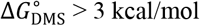 (black in Fig. 5A; fig. S13). All other canonically paired WCF nucleotides exhibit 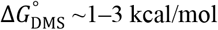, consistent with our earlier measurements in the fourU and TAR RNAs. Similarly, nucleotides participating in local intra-helical non-canonical pairs exhibit 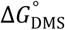 values between 0 and 2 kcal/mol. As an exception, several Hoogsteen-paired adenosines exhibit strongly negative free energies, consistent with the N1 being oriented into solution by this pairing geometry (for example, A207 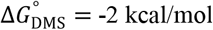) (fig. S16). Unpaired nucleotides exhibit 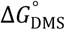 of 0.2 to 1.0 kcal/mol, also congruent with ranges observed in fourU and TAR RNAs. Thus, the energetic scale of local nucleotide fluctuations remains largely comparable across simple hairpins and complex three-dimensional RNA folds.

**Figure 5.**
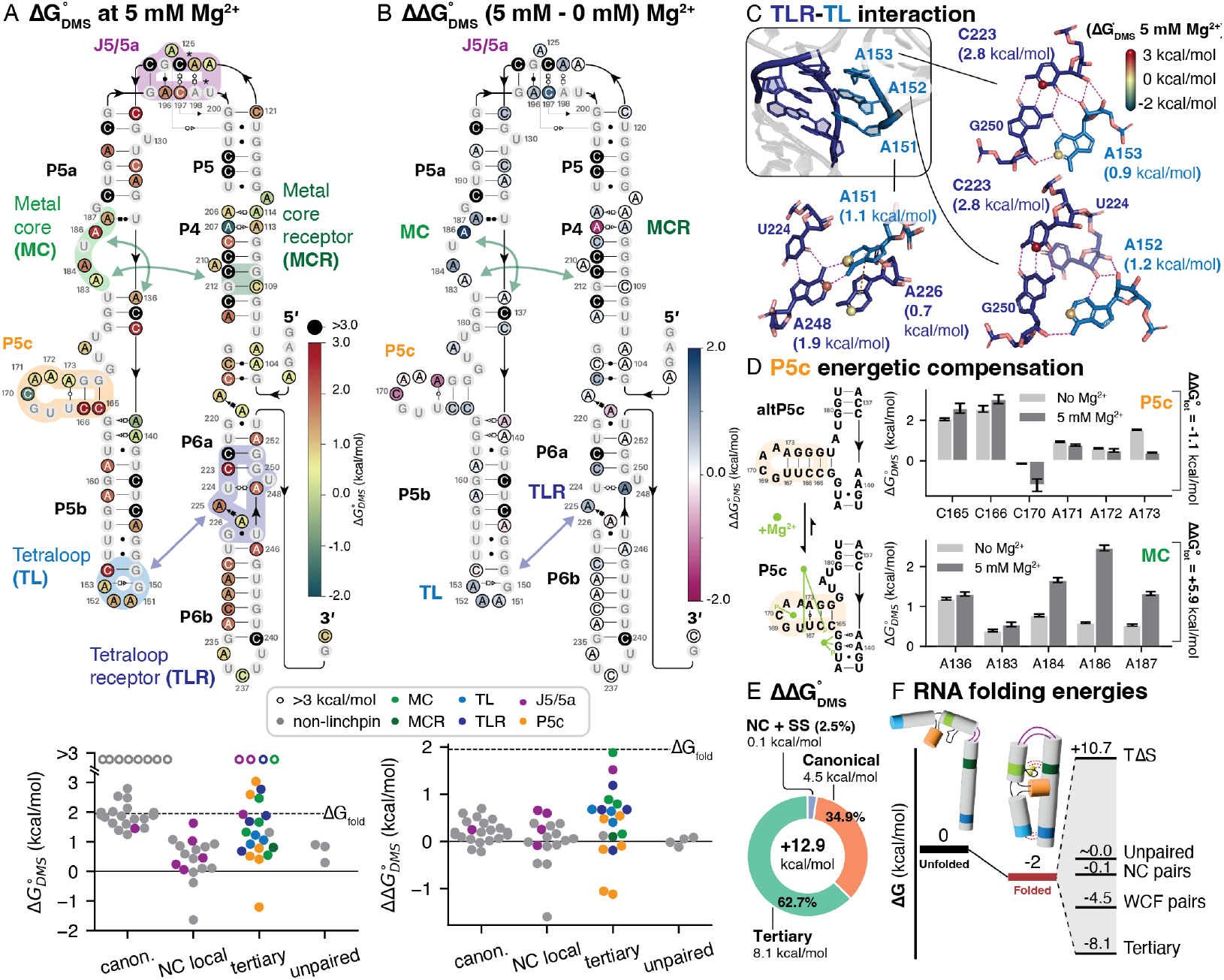
PRIME-DMS nucleotide free energies reveal energetic coupling between secondary and tertiary folding in the P4-P6 domain of the Tetrahymena ribozyme. (**A**) Annotated secondary structure of P4-P6 with base pairs classified using Leontis-Westhof geometry. The tetraloop-tetraloop receptor (TL-TLR), J5/5a junction, P5c, and metal core-metal core receptor (MC-MCR) regions are highlighted. Nucleotides are colored according the 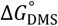 values at 5 mM Mg^2+^. Swarm plots below show distributions grouped by canonical Watson-Crick-Franklin (WCF), local non-canonical (NC local), tertiary interactions, and unpaired sites. Highest reactive sites (C125 and A198; asterisks) were used to estimate *k*_add_ for C and A, respectively. (**B**) Mg^2+^-dependent changes in nucleotide energetics (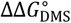, 5 mM - 0 mM Mg^2+^) with corresponding swarm plots grouped as in (A). (**C**) 3D view of the TL-TLR interface (matching color scheme as in (A)) with 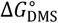 (5 mM Mg^2+^) mapped onto spheres at N1 of A and N3 of C, color-coded by energy; key hydrogen bonds are shown in magenta. (**D**) P5c subdomain comparison. (left) Secondary structures of the native and alternative P5c conformations (*52*). Green circles indicate Mg^2+^ ions observed in the crystal structure (PDB 1GID), green lines indicate Mg^2+^ coordination to phosphate oxygens. (right) Site-resolved 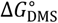 with and without Mg^2+^ for P5c and MC regions. Summed 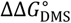 values are indicated on the right, revealing net destabilization in P5c and net stabilization in the MC. (E) Breakdown of 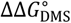 by interaction type, with the total summed value indicated in the center. (F) Schematic illustrates energetic compensation: folding incurs an entropic penalty but is offset by stabilizing secondary and tertiary interactions, yielding the observed net global energetics.

P4-P6 also features multiple long-range tertiary pairing interactions, the energetics of which have never before been resolved at nucleotide resolution. The P4-P6 global fold is defined by the tetraloop/tetraloop receptor (TL/TLR) interaction between A151-A153 and C223, A225/226, A248 (*24*) (Fig. 5C). A151-A153 form a diverse network of hydrogen bonds to the TLR involving base-base, base-sugar, and sugar-sugar contacts (Fig. 5C). Each TL adenosine exhibits similar 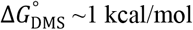, indicating a low cost of individual opening. By comparison, TLR nucleotides possess 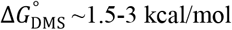 (Fig. 5A), consistent with an asymmetric energetic distribution and enhanced TL dynamics inferred from prior NMR studies (*47, 48*). Another major tertiary motif is the metal core/metal core receptor (MC/MCR), which involves extensive local stacking and non-canonical interactions within the MC, and long-range tertiary pairs formed between the MC to the MCR. MC nucleotides exhibit a range of 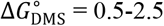 kcal/mol, with the linchpin nucleotide A186 demonstrating the greatest 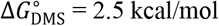.

Nucleotides A183 and A184, which form long range base-triple pairs to the MCR, demonstrate 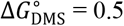 and 1.6 kcal/mol, respectively. The J5/J5a hinge also features several important local tertiary pairing interactions that stabilize the sharp turn between P5 and P5a, which demonstrate 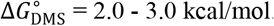. Because global 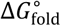 is expected to be ∼2 kcal/mol under our probing conditions (*46*), these data thus indicate a tiered hierarchy of tertiary fluctuations. Nucleotides featuring 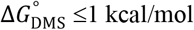, including A151-A153 and select nucleotides in the MC/MCR, likely undergo local opening fluctuations within the intact 3D fold. By contrast, nucleotides in the MC/MCR and hinge that feature 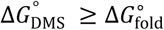 are likely gated by global tertiary unfolding (Fig. 5A, above dashed line). Notably, these ‘high energy’ MC/MCR and hinge nucleotides correspond to previously identified ‘linchpin’ interactions essential to P4-P6 folding (*24, 46, 49, 50*), whereas the TL is more tolerant to individual mutation (*51*).

To assess the dependency of nucleotide energetics on tertiary structure, we repeated PRIME measurements on P4-P6 folded in the absence of Mg^2+^, where it lacks tertiary structure, and calculated the 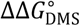 between +Mg^2+^ and -Mg^2+^ conditions (Fig. 5B). As expected, the largest 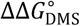 values localize to known tertiary contacts. 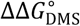 within the TL/TLR were 0.4-0.7, consistent with a more dynamic interaction. Interactions in the MC and hinge show 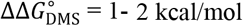, close in magnitude to the global folding free energy. Surprisingly, many canonical WCF pairs also show small increases in stability (average 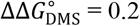; Wilcoxon p-value = 0.001; Fig. 5B). These changes mostly localize to junction-closing pairs, but diverse internal pairs also demonstrate modest increases in stability (e.g. C229 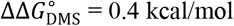). Together, these results indicate that Mg^2+^-dependent stabilization of tertiary interactions propagates beyond discrete contact sites, subtly reinforcing secondary structure throughout the molecule and contributing to overall fold stability.

Significantly, PRIME also directly resolves energetic tradeoffs within the P5c domain, which has long served as an exemplar of secondary-tertiary compensation but only indirectly measured via mutagenesis (*52, 53*). In the absence of Mg^2+^, P5c adopts an alternative secondary structure (altP5c) with two additional base pairs, which must be disrupted to enable metal core formation upon Mg^2+^ binding (Fig. 5D). PRIME captures this compensation as a net destabilization of the P5c region upon Mg^2+^ binding (total 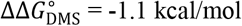; Fig. 5D). Notably, this local energetic penalty is more than offset by strong Mg^2+^dependent stabilization of the adjacent metal core 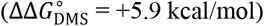, where every adenosine in the region becomes more stable (Fig. 5D). Thus, formation of the Mg^2+^-stabilized MC enhances stability of nucleotides throughout the P5abc junction, compensating for the loss of several pairs compared to altP5c.

We finally explored how nucleotide-level energetics contribute to the global energetics of Mg^2+^-dependent tertiary folding. Summing 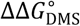 across all P4-P6 nucleotides indicates tertiary folding yields +12.9 kcal/mol of local nucleotide stabilization (Fig. 5E). Previous studies of toy models have suggested that, under comparable buffer conditions to those used here, tertiary folding of P4-P6-like molecules incurs a free energy penalty of 6-11 kcal/mol, predominantly due to loss of global conformational entropy (*54*), remarkably close to the 10.7 kcal/mol value needed to counteract the 12.9 kcal/mol net local stabilization measured by PRIME to arrive at a global 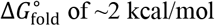 measured by smFRET (*46*) (Fig. 5F). Significantly, while 8.1 kcal/mol of this stabilization (63%) derives from classical tertiary interactions, the remaining 4.5 kcal/mol (35%) derives from increased stability of other interactions, including WCF pairs (Fig. 5F). Thus, our data support a model in which secondary-structure stabilization is a key contributor to Mg^2+^-dependent folding.

## Discussion

RNA folding energetics govern the shapes, populations and dynamics of RNA structures and their interactions with cellular factors, providing the physico-chemical foundation for understanding RNA functions (*4*). However, most knowledge about RNA thermodynamics is derived from measurements of global RNA stability (*55*). NMR can provide direct nucleotide-resolution measurements, but is limited by RNA size and throughput and generally cannot be performed in complex biological contexts. Accumulating evidence has shown that chemical probing reactivities correlate with thermodynamic folding free energies (*56-59*). PRIME extends this observation, providing a biophysical framework to quantitatively deconvolve chemical probing data into intrinsic nucleotide energetics comparable to NMR. Similar to HDX for proteins (*11, 21*), PRIME is able to go beyond static structures, uncovering hidden features of RNA folding landscapes such as coupling between secondary and tertiary structure that have remained challenging to access with traditional biophysical methods.

Application of PRIME to three model RNA systems of varied complexity uncovered that RNA nucleotides ubiquitously undergo low energy (< 3 kcal/mol) structural fluctuations at 25°C. Thus, even ‘strong’ WCF RNA base pairs are much more dynamic than commonly assumed, corroborating early tritium exchange experiments that have remained controversial (*8*). Specifically, an energy of 2 kcal/mol represents a 3.3% probability that a paired base will be accessible at 25°C, facilitating rearrangement via strand displacement (*60*) or recognition by proteins (*4*) or the ribosome (*61*), among other functionally important interactions. PRIME also reveals that local non-canonical and long-range tertiary pairs have a wide-range of stabilities, with, for example, the P4-P6 TL nucleotides likely undergoing local opening dynamics within the 3D fold, whereas other tertiary interactions are likely gated by global folding.

PRIME has several practical and conceptual limitations that define immediate opportunities for expansion. Estimates of 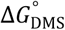 depend on accurate determination of base-specific *k*_add_. This approximation may overlook context-dependent biases in RT detection efficiency of chemical probing adducts. In its current formulation, *k*_add_ is estimated from nucleotides expected to be completely unprotected. Residual structure may lead to underestimation of *k*_add_, and correspondingly inflated 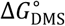 values. We focused here on nonlinear fitting of full time-course data to establish rigorous validation, but extension of PRIME to endpoint measurements is readily achievable. Despite these limitations, PRIME can be implemented with simple additions to existing chemical probing protocols, is highly scalable, free of size constraints, and substantially simpler to implement than NMR (*62*).

We envision several immediate extensions and applications of PRIME. Extension to other chemical probes such as 2′-OH acylation (*63*) promises to expand our knowledge of other types of RNA fluctuations. PRIME energies can also provide a principled foundation for correcting well-known energetic deficiencies in current RNA folding algorithms (*64*), as well as improving molecular dynamics force fields (*65*). PRIME can also be used to more rigorously quantify the structural impact of natural RNA modifications, and provide new routes to understanding their functional roles in biology (*66*). When leveraged at scale, and eventually extended to cellular probing, PRIME will be able to generate much needed biophysically rigorous datasets to power AI-driven RNA modeling and design (*67*). In sum, PRIME establishes a new foundation for nucleotide-resolution RNA biophysics (*68*) with broad potential to improve understanding of biological mechanisms and enable RNA bioengineering.

## Materials and Methods

### RNA design

WT and mutant RNA sequences for *Salmonella enterica* agsA fourU thermometer (*28*), HIV-1 TAR RNA (*35, 37, 38*), and *Tetrahymena thermophilus* ribozyme P4-P6 domain (*24*) were obtained from previous publications as cited. Following published protocols (*69*), each construct was extended to include a 19-20 nt 5′-primer binding site and 3′ reverse transcription (RT) binding site flanking the target RNA sequence. For the fourU thermometer and TAR constructs, synthetic sequences were designed using NUPACK (*70*) to minimize spurious base-pairing with the native sequence (**Supplementary Note 5**). For the P4-P6 construct, flanking sequences from (*49*) were slightly modified to add a 6 nt spacer between primer binding sites and the target sequence (**Supplementary Note 5**). All constructs used are listed and annotated in **table S1**.

### RNA synthesis and purification

Large-scale synthesis and purification of RNA was based on a previously published protocol (*71*) with modifications. DNA templates were ordered as custom gBlocks (IDT) and amplified by PCR using Q5 DNA Polymerase (NEB Cat. No. M0491L) and relevant primers (**table S2**). Large-scale *in vitro* transcription reactions (1-3 mL) were performed (40 mM Tris-Cl, pH 8, 14 mM MgCl_2_, 2 mM spermidine, 2.5 ng/µL DNA template, 100 µg/µL homemade T7 RNAP). Following transcription, RNA was extracted using 1 vol. phenol:chloroform:isoamylalcohol 25:24:1 (Sigma-Aldrich Cat. No. 516726-1SET) and 1 vol. chloroform (Fisher Scientific Cat. No. 423555000). At each extraction step, the transcription mix was mixed vigorously then centrifuged for 6-8 mins. at 21,130 rcf before the aqueous layer was carefully extracted for the next step. To remove salts and some unincorporated nucleotides, the RNA was further purified by gel-filtration. Extracted transcription mix was buffer-exchanged into gel-filtration buffer (15 mM K_x_(PO_4_)_y_ pH 6.5, 100 mM KCl) using Amicon Ultra-15 MWCO 3 kDa (Millipore-Sigma Cat. No. UFC900308) or Vivaspin 15R MWCO 2 kDa (Vivaproducts Cat. No. VS15RH91) columns. Purification was then carried out on an ÄKTAxpress FPLC system using a Superdex 75 16/600 column (Cytiva Cat. No. 28989333). Resulting chromatograms generally showed 1 fast-running peak for residual NTPs and 1-2 slower peaks containing the target RNA. As confirmed on a denaturing analytical gel, peak fractions containing the target RNA were selected and pooled in ∼8-10 mL. Next, RNA was buffer-exchanged into double-distilled water (ddH_2_O) through dialysis or concentrator columns. Dialysis was performed in a Spectra/Por 8 kDa MWCO dialysis bag (Spectrum Cat. No. 132660) against 2 L ddH_2_O for 12 hrs, changing water every 4 hrs. The final sample was then concentrated down to 1 mL using Amicon Ultra-15 MWCO 3 kDa (Millipore-Sigma Cat. No. UFC900308) or Vivaspin 15R MWCO 2 kDa (Vivaproducts Cat. No. VS15RH91). If dialysis was not performed, then 3 additional buffer-exchanges against 15 mL ddH2O was performed on the columns before final concentration. RNA was finally quantified using Nanodrop and verified for purity via analytical denaturing PAGE.

### Buffer preparation

Dimethyl sulfate (DMS) decomposes into sulfuric acid (**fig. S7**), resulting in reductions in pH if the solution is not sufficiently buffered (**fig. S2**) (*29*). We therefore optimized buffer concentrations to maintain pH throughout a time-course reaction and chose appropriate buffers based on the required pH for each RNA construct used (**fig. S2C**). The composition of the buffers are as follow: fourU thermometer (150 mM bis-tris pH 6.5, 15 mM K_x_(HPO_4_)_y_, 25 mM KCl), TAR (150 mM bis-tris pH 6.4, 15 mM Na_3_PO_4_, 25 mM NaCl, 0.1 mM EDTA), P4-P6 (200 mM bicine pH 8.0, 100 mM KCl, 0 mM or 5 mM MgCl_2_). Separate buffers were prepared for use at different folding temperatures to account for temperature-dependence of pH. *d*(*pK*_*a*_)/*dT* values of -0.017 (*72*) for bis-tris buffers, and -0.018 (*73*) for the bicine buffer were used to calculate expected pH at room temp (21°C - 22°C) (**Supplementary Note 6**).

### Time-course chemical probing

Each time-course experiment (6 time points) was conducted at a specific target temperature. Experiments were conducted on a metal heat block, and reaction temperatures were calibrated prior to each experiment using a glass thermometer inserted into a dummy tube containing ddH_2_O and equilibrated in the same heat block. For each experiment, 84 µL 1.375 µM RNA in ddH_2_O was denatured at 95°C for 3 mins then snap-cooled on ice for at least 5 mins. Folding was initiated by adding 21 µL of 5× folding buffer (as detailed above) and equilibrated at the target temperature either for 20 mins (target temp 12°C-60°C), or 5 mins (target temp 60°C to 80°C). Tubes were set up for the experimental condition (+) with 9.6 µL 1.5% (v/v) DMS diluted in 100% ethanol (Fisher Scientific Cat. No. 04-355-720) and control condition (™) with 1.6 µL 100% ethanol. After temperature equilibration, the probing reaction was initiated by adding 86.4 µL 1.1 µM RNA in folding buffer to the (+) reaction tube, while 14.4 µL 1.1 µM RNA is added to the (™) reaction tube, yielding final probing reaction conditions of 1 µM RNA and 0.15% DMS (v/v) in 1× folding buffer. Both tubes are held at a target temperature for the duration of a reaction. To capture time courses, aliquots of probing reaction were quenched at indicated timepoints by withdrawing 15 µL of the sample and vigorously mixing into 250 µL quench mix (Final conc.: 4.5 M β-mercaptoethanol (Millipore-Sigma Cat. No. 444203), 300 mM NaOAc, pH 5.2) using a vortex and immediately transferred to ice. The same process is applied to the untreated (-) sample at the longest time point. After all time points were taken, quenched RNA was ethanol precipitated by adding and mixing 26.5 µL 3 M sodium acetate, 0.6 µL 0.1 mg/mL GlycoBlue (ThermoFisher Cat. No. AM9515) and 662.5 µL 100% ethanol. After chilling for 30 mins at -80°C, RNA pellets were recovered by centrifuging at 21,130 rcf for 30 mins. Pellets were washed twice with 1 mL 75% ethanol, air-dried, and resuspended in 20 µL nuclease-free ddH_2_O. Temperature and replicate numbers for all time-course probing experiments are listed in **table S3**. We note that endpoint measurements at the longest time point (8 h at 23°C) demonstrated robust agreement of fitted parameters across independent replicates (**fig. S15**). Based on this reproducibility, subsequent analysis of the P4-P6 domain was performed using a single full time-course at 23°C for each Mg^2+^ condition.

### Reverse transcription

Reverse transcription was performed using MarathonRT (Kerafast Cat. No. EYU007) following a modified protocol based on Mitchell et al (*30*). To set up a 20 µL RT reaction, 5 µL RNA (∼1 µM) was combined with 1 µL 10 mM dNTPs, 0.5 µL 10 µM RT primer for the RNA under study (**table S4**), and 0.3 µL ddH_2_O. The mixture was denatured at 95°C for 3 minutes and annealed at 65°C for 5 minutes, then chilled on ice. RT buffer (12.2 µL of 5× MarathonRT buffer containing 50 mM Tris-HCl, pH 8.3, 200 mM KCl, 5 mM DTT, 1 mM MnCl_2_, 20% glycerol) was added, followed by 1 µL MarathonRT enzyme. Reactions were incubated at 42°C for 3 hours and heat-inactivated at 95°C for 1 minute. RNA was hydrolyzed by adding 1.25 µL of 4 M NaOH and incubating at 95°C for 5 minutes, followed by neutralization with 2.5 µL of 1 M HCl. The cDNA was purified using 1.8× volume SPRI beads (Beckman Coulter AmpureXP Cat. No. A63881 or Cytiva Sera-Mag™ Select, Cat No. 29343052) and eluted in 30 µL nuclease-free water.

### Library preparation and sequencing

cDNA libraries were prepared using a two-step PCR amplification strategy using primers listed in **table S4**. For all PCR reactions, we used Q5 High-Fidelity DNA polymerase (NEB Cat. No. M0491L) and followed standard manufacturer protocol, scaled down to 10 µL. In PCR1, 1 µL of cDNA was amplified using the indicated PCR1 primers with the following cycling conditions: 98°C for 30 s; 15 cycles of [98°C for 10 s, 68°C for 30 s, 72°C for 20 s]; final extension at 72°C for 2 min. PCR1 products were purified with a 1.8× SPRI bead cleanup and eluted in 30 µL ddH2O.

Then, 1 ng of purified PCR1 product was used as input for PCR2 using a set of unique dual-indexed 8-bp PCR2 primers: 98°C for 30 s; 10 cycles of [98°C for 10 s, 69°C for 30 s, 72°C for 20 s]; final extension at 72°C for 2 min. An Echo 525 Acoustic Liquid Handler (Beckman-Coulter) was used to transfer unique index primers into separate wells. PCR2 products were purified as PCR1. Final libraries were quantified using the Qubit dsDNA HS Assay Kit (ThermoFisher Cat. No. Q32851) or Quant-iT dsDNA HS Kit (ThermoFisher Cat. No. Q33120).

Pooled libraries were quantified using a Qubit dsDNA HS assay and quality-controlled using a Bioanalyzer DNA HS or DNA 1000 assay. Libraries were then sequenced on an Illumina MiSeq (2 × 75 bp, v3 chemistry, 15-16 pM loading concentration) or NextSeq 1000/2000 (2 × 150 bp, P1 kit, 400-450 pM loading concentration) with 10-15% PhiX spike-in.

### Mutational rate analysis

Mutational profiling of sequencing FASTQ files was performed using ShapeMapper v2.3 (*62*). Both DMS-treated and untreated samples were processed identically using the --modified flag without any background subtraction or normalization.

### Command

`shapemapper --name $name --target $target_seq --modified --R1 $treated_r1 --R2 $treated_r2 --dms --amplicon`

Here, `$` denotes variable name in bash, $name denotes a sample identifier, `$target_seq` the reference RNA sequence in FASTA format, and `$treated_r1/$treated_r2` the paired-end FASTQ files. The `--dms` flag enables DMS-specific mutation calling, while `--amplicon` specifies targeted amplicon sequencing.

Nucleotide reactivities were computed from per-position mutation rates inferred from reverse-transcription misincorporations, defined as the number of observed mutations divided by the effective read depth after quality filtering and adduct localization. Reactivities, *r*_*j*_, in text refers to the `Modified_rate` output column reported by ShapeMapper and used directly for time-course fitting.

ShapeMapper execution was orchestrated within the *<u>nerd</u>* framework and is available in the *<u>nerd</u>* GitHub repository (see `pipeline/plugins/mutcount/shapemapper.py`, function `command`).

### Time-course fitting of probing reactivities

For each RNA studied, and each temperature condition, time course mutational rates, or reactivities, for each nucleotide *j, r*_*j*_, were fit according to the closed form solution to the chemical kinetics model of the probing reaction Eq. (1) (see **Supplementary Note 1** for full derivation). All non-linear least-squares regression were performed using the *lmfit* Python library(*74*), generating 2 fitted parameters, *k*_deg_ and *k*_obs,j_, where

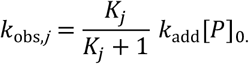

For each RNA and temperature condition, we first performed independent time-course fits at every adenosine (A) and cytidine (C) position. The resulting datasets were then used to determine a single global value of *k*_deg_, obtained by fitting this constant across all positions simultaneously. Using this globally estimated *k*_deg_, the time courses were then refitted to obtain the final *k*_obs,j_ values. See **Supplementary Note 1** and **Data, code, and materials availability** for the Python code implementation of this fitting procedure.

### Global adduction rate (*k*_add_) estimation

For each RNA studied, we estimated adduction rates (*k*_add_) for adenosine (A) and cytidine (C) bases using fitted *k*_obs,*j*_ values according to the approaches in **Supplementary Note 2**. We used Strategy 1 for the fourU RNA, Strategy 2 for the HIV TAR RNA, and Strategy 3 for the P4-P6 RNA. For Strategies 2, to obtain *k*_add,A_ for a given RNA, we first averaged ln;k_obs,*j*_) across all A positions and independent replicates at each temperature (>60°C). The resulting mean ln;k_obs,A_) values at each temperature >60°C were then plotted as a function of reciprocal temperature (1/T). Linear regression using Eq. (16) in **Supplementary Note 2**, was used to fit *E*_*a*,A_ and prefactor *A*_A_, using the known value of [*P*]_0_. An identical procedure was performed for cytosines. For Strategy 3, the maximum ln(*k*_obs,*j*_) for A positions and C positions are used as ln(*k*_add,A_) and ln(*k*_add,C_), respectively.

Linear regression fits were carried out using the LinearModel class from the *lmfit* Python library (*74*). See **Supplementary Note 2** and **Data, code, and materials availability** for the Python code implementation of this fitting procedure.

### Arrhenius analysis of kinetic rates

Temperature dependence of extracted kinetic rates was analyzed using the Arrhenius equation, *k*(*T*) = *A* exp(™E_*a*_/*RT*). Rate constants were linearized as

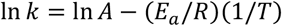

and plotted as a function of reciprocal temperature (1/T, in K^-1^). Linear regression was used to extract activation energies *E*_*a*_ from the slope and prefactors *A* from the intercept. Fits were performed using the LinearModel class from the *lmfit* Python library (*74*).

Calculation of per-nucleotide 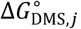 from *k*_obs_

For each RNA, values of *K*_*j*_ at each temperature studied were extracted from *k*_obs,*j*_ values following the approach in **Supplementary Note 2**, where

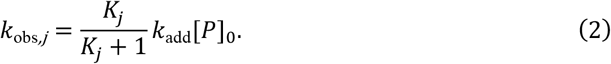

Strategy 1 was used for the fourU system, Strategy 2 was used for the TAR system, and Strategy 3 was used for the P4-P6 system.

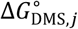 values were calculated according to **Supplementary Note 3**. For comparisons against reference energies, 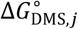 was obtained at 25°C for the fourU thermometer and for HIV-1 TAR; 23°C for P4-P6 to match conditions of the original studies.

### Uncertainty sources and error propagation

Uncertainties arise from two primary sources: (i) non-linear fitting of time-course data to obtain ln *k*_obs,j_ with uncertainty *σ*_1,*j*_, and (ii) estimation of ln(*k*_add_(*T*) [*P*]_0_) from high-temperature extrapolation with uncertainty σ_2_. These errors are propagated using first-order error propagation (**Supplementary Note 4)**, yielding

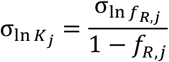

where *f*_R,*j*_ = *K*_*j*_/(1 + *K*_*j*_). Finally, uncertainties in nucleotide-level energetics are computed from

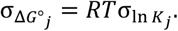

For independent biological replicates, uncertainties are combined by summing the squared replicate errors and dividing the square root by the number of replicates, 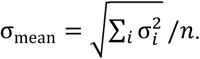.

## See Supplementary Note 4 for full derivation

### Nucleotide-resolution energy benchmarking

NMR-based energies for the fourU system were taken from Schwalbe et al. (*28*). These are derived from imino-proton exchange experiments, which track H-donors within a base-pair at guanosines and uridines. Nearest-neighbor (NN) model-based energies were calculated as detailed below. Transformation of chemical probing reactivities into pseudo-energies was performed as

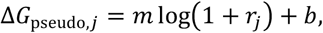

where we used *m* = 1.1 and *b* = ™0.6 (*75*). Reactivities *r*_*j*_ were estimated by evaluating the fitted time-course model at *t* = 10^6^ s, which provides the best estimate of the true reaction endpoint. Agreement between datasets was assessed by least-squares linear regression. Comparable results were obtained when using reactivities at 15 min or raw data points, with no improvement in R^2^ (data not shown). See **Data, code, and materials availability** for the Python code implementation for reactivity processing and correlation fitting.

### Nearest-neighbor (NN) energy analyses

The NN model describes RNA folding stability as the sum of local energetic contributions that are minimized to determine the most stable secondary structure. In this framework, the fundamental energetic unit is a nearest-neighbor stack consisting of two adjacent base pairs, rather than an individual nucleotide, complicating direct comparison with nucleotide-resolved experimental measurements.

To enable such comparisons, we implemented an NN-based “partition function approach” using the RNAstructure program (*32*). For each nucleotide, the position was forced to be unpaired during partition-function calculation. The free energy difference between the constrained (forced-unpaired) and unconstrained ensembles represents the NN energetic cost of rendering that nucleotide single-stranded within the full structural ensemble. Δ*G*° values were extracted using Turner 2004 NN parameter set (*76*). See **Data, code, and materials availability** for the Python code implementation for calculation of NN-based energies.

### Independent measurement of DMS kinetics with 1D NMR

For DMS degradation measurements, samples (750 µL) were prepared in 5 mm Wilmad NMR tubes containing 150 µL of 5× fourU pH 6.5 folding buffer (see **Time-course chemical probing**), 75 µL D_2_O, and 450 µL ddH_2_O. This reaction mix was pre-equilibrated in the NMR spectrometer (Bruker Avance III 600 MHz A600) for ∼ 20 mins. In parallel, 100 µL of 1.5% DMS (diluted in d6-EtOH) was pre-incubated separately at the target temperature using a calibrated metal heat block. To initiate the reaction, the pre-incubated NMR tube containing 750 µL 1× folding buffer, 10% D_2_O was ejected from the machine and 75 µL 1.5% DMS was added directly to the tube. Upon recapping, the tube was carefully rotated three times to mix, and the true reaction start time noted. The tube was then re-inserted into the spectrometer, where routine instrument preparation (locking, shimming, and receiver-gain calibration) was performed. Scan parameters are listed in **table S5**, and the Bruker TopSpin *multi_zgvd* program was initiated immediately afterward.

DMS and byproduct peaks were identified using previously published assignments (*27*). Time-resolved DMS signal loss was then fit using a three-parameter exponential decay model:

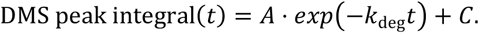

For NTP adduction rates, samples were prepared as described above with the addition of 5 mM ATP or CTP. The same steps were followed above except some data was collected using a Brucker Neo 600 MHz HFCN600 instrument. Resonances corresponding to NTP peaks were identified using reference spectra from the Human Metabolome Database (*77*) (ATP: HMDB0000538, CTP: HMDB0000082). For ATP, the C8 (∼8.0-8.3 ppm) and C1’ (∼5.9-6.2 ppm) resonances exhibited a pronounced upfield shift. In CTP, the C6 (∼7.8-8.1 ppm) and C5 (∼6.1-6.3 ppm) positions showed similar behavior. Because these positions are electronically coupled within the nucleobase ring system, their upfield displacement is consistent with progressive NTP methylation. These resonances, together with the DMS peak decay, were used to extract *k*_add_ values. The kinetics were modeled using a system of ordinary differential equations (ODEs) describing two coupled processes: methylation of native signal (1) and degradation of the reactive species (2).

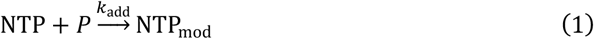

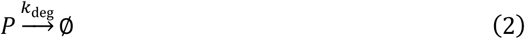

which can be modeled by:

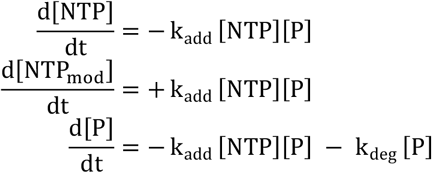

Using *solve_ivp* from Scipy (*78*), these ODEs were numerically solved for the parameters *k*_add_ and *k*_deg_ using initial conditions, *k*_add_ = 0.03, *k*_deg_ estimated from independent DMS degradation experiments, [NTP_U(>_] = 0 mM, [*P*] = 15.64 mM. Initial concentrations of NTP, [NTP], were independently measured using A260 measurements from Thermofisher NanoDrop OneC (Cat. No. ND-ONE-W) rather than using nominal concentrations: ATP 8.00 mM, CTP 7.36 mM. Kinetic parameters were fitted using scipy.optimize functions in Python. See **Data, code, and materials availability** for the Python code implementation of this fitting procedure.

## Acknowledgments

We would like to acknowledge Susan Marqusee and Charles Brooks for early suggestions to consider applying the HDX framework to chemical probing, and Chaitan Khosla for suggesting that chemical probing data could be used to uncover RNA folding energetics. We thank Aria Coraor Adam Silverman for early work on deriving the PRIME mathematical framework, Kayd Bhagat for code review, and Molly Evans and Khoa Dao for performing early experiments. We thank Gabe Rocklin, Dan Herschlag, Joe Yesselman, John Shin, Rhiju Das, Hashim Al-Hashimi, Yeongjoon Lee, and members of the Lucks lab for helpful conversations. Any opinions, findings and conclusions or recommendations expressed in this material are those of the author(s) and do not necessarily reflect the views of the National Science Foundation, the National Institutes of Health or the Simons Foundation.

## Author contributions

Conceptualization: EKC, JBL, AMM

Methodology: EKC, RB

Investigation: EKC, RB, DHM, JBL, AMM

Formalization: EKC

Visualization: EKC, JBL, AMM

Funding acquisition: JBL, AMM

Project administration: EKC, JBL, AMM

Software: EKC

Supervision: EKC, JBL, AMM

Writing – original draft: EKC, JBL, AMM

Writing – review & editing: EKC, DHM, JBL, AMM

## Data, code, and materials availability

The core PRIME data analysis framework (nerd) is available at (https://github.com/LucksLab/nerd). All source data supporting these findings, including SRA accession numbers, together with the analysis code corresponding to each figure, are archived in a Zenodo repository (https://doi.org/10.5281/zenodo.18398129), which includes a README describing the organization of the data and analysis workflows. Specific functions used in the analyses are cited in the Methods and Supplementary Notes and are available within the Zenodo archive.

## Supplementary Materials

Materials and Methods

Supplementary Text

Figs. S1 to S16

Tables S1 to S7

References (*1*–*65*)

