## Supplementary Information for "Ubiquitous low-energy RNA fluctuations and energetic coupling measured by chemical probing"

**The PDF file includes:**

Figs. S1 to S16
Supplementary Text
Tables S1 to S6
References

### Supplementary Figures

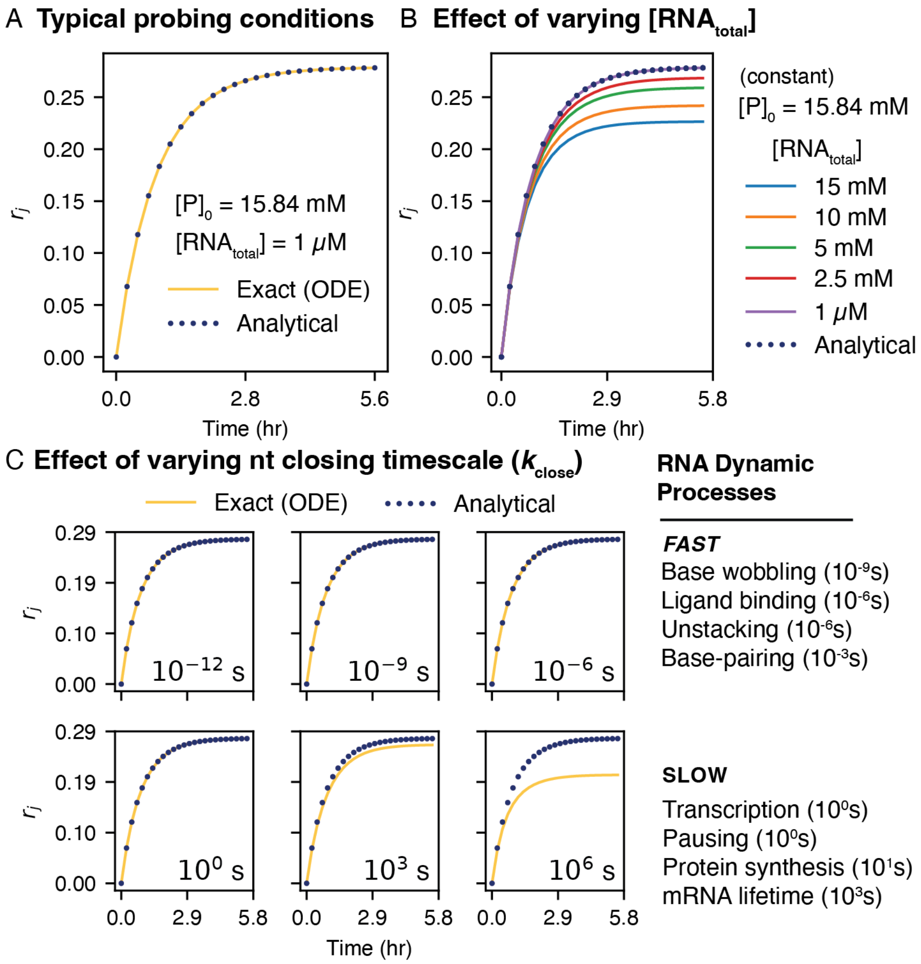


**Figure S1. The analytical solution to the chemical probing kinetic equations matches the exact solution under typical probing conditions and RNA fluctuation times.** (**A**) The analytical solution for reactivities (Supplementary Note 1) matches ODE simulations: $[RNA_{\text{total}}]$ = 1 μM, $[P]_0$ = 15.84 mM, with probe kinetic rates estimated using unpaired A18 in a representative WT fourU RNA time-course at 25°C, $k_{\text{add}}$ = 0.018 $M^{-1}s^{-1}$, $k_{\text{deg}}$ = 0.0029 $s^{-1}$, and modestly fast open/close rates, $k_{\text{open}}$ = 100 $s^{-1}$, $k_{\text{close}}$ = 200 $s^{-1}$. (**B**) To test the validity of the analytical solution as probe depletion due to RNA modification becomes non-negligible, we varied the total RNA concentration $[\text{RNA}_{\text{total}}]$ while holding the initial DMS concentration $[P]_0$ constant. All other parameters as in (A). The analytical and exact (ODE) solutions agree over a broad range of RNA concentrations but begin to deviate as $[\text{RNA}_{\text{total}}]$ approaches the DMS concentration (15.84 mM). ODE code implementation is provided in the Zenodo deposit (see **Data, code, and materials availability**). (**C**) Comparison of the analytical solution and ODE simulations for different values of $k_{\text{close}}$ (indicated) shows the analytical solution remains accurate for nucleotide fluctuations >>1 s timescales. All other parameters are held constant as in (A).

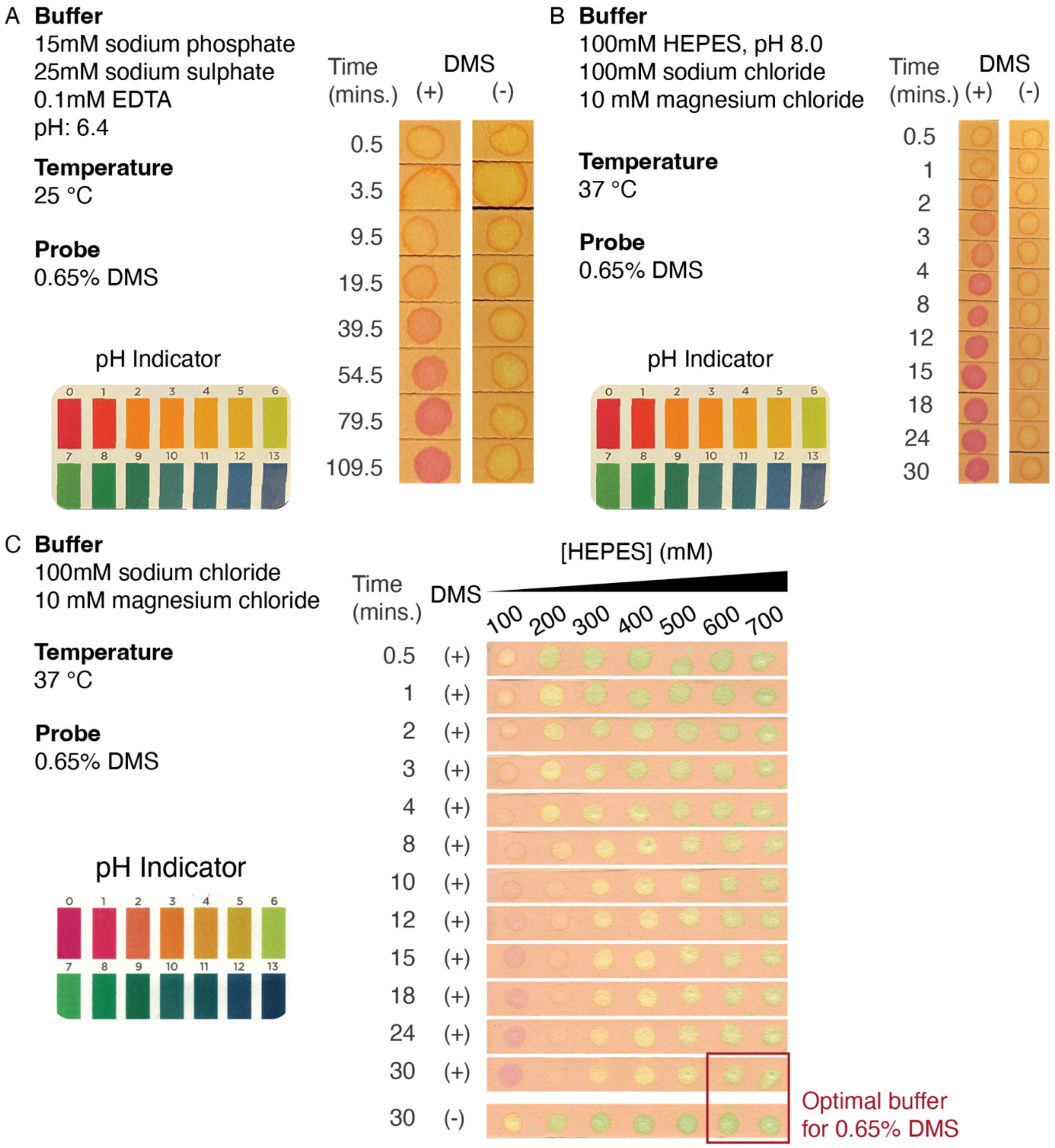


**Figure S2. Optimizing probing reaction conditions to buffer against DMS hydrolysis acidification.** (**A, B**) Typical buffers used for DMS probing acidify during the probe reaction. Photographs are of pH strips blotted with probing reactions either treated with (+) the DMS concentration or an ethanol control (-). (**C**) Titration of HEPES concentration identified an optimal buffering range of 600–700 mM, which maintained pH ≈ 8 for ≥30 min (reaction completion) at 37°C in the presence of 0.65% DMS. These measurements guided the selection of the final reaction conditions used throughout this study (150 mM HEPES, 0.15% DMS).

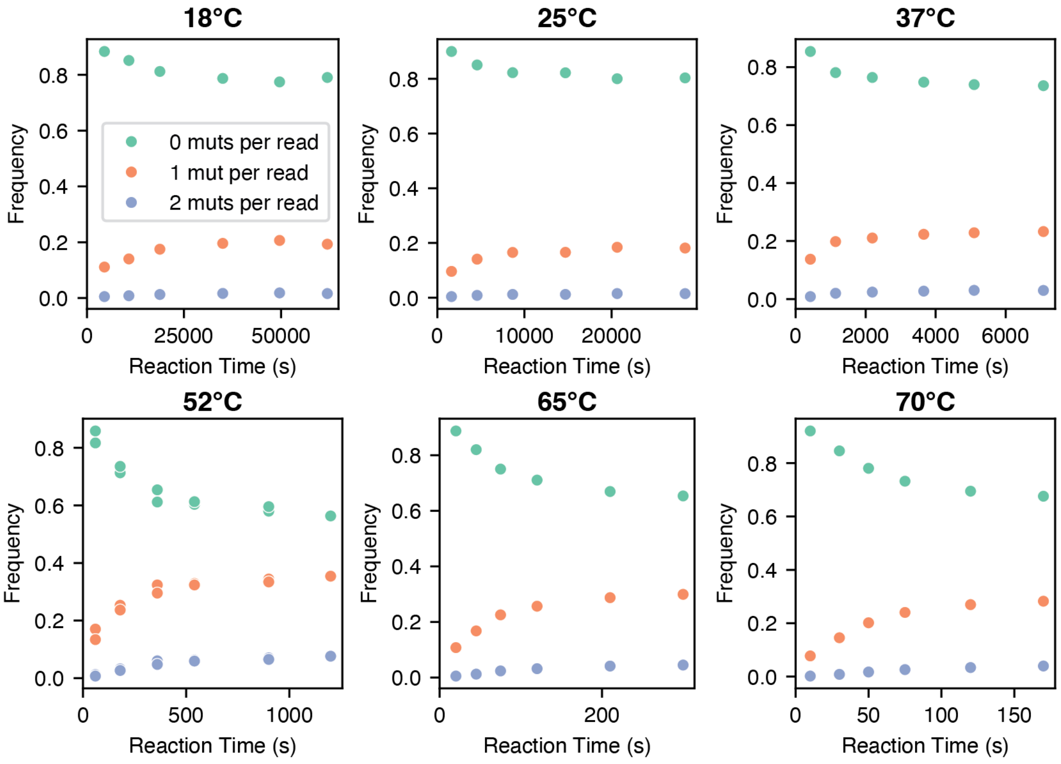


**Figure S3. Assessment of single-hit regime.** Representative DMS-MaP time-course profiles grouped by number of mutations per read for the WT fourU RNA collected at representative replicates at 18°C, 25°C, 37°C, 52°C, 65°C, and 70°C. Reactions were performed at 15.84 mM (0.15% v/v DMS) with 1 µM RNA, and mutations were mapped and calculated using ShapeMapper2.3. Across a broad range of temperatures and reaction times, 18-35% of reads contained a single mutation, whereas only 1-4% contained two mutations, distributed across the 80-nt RNA, confirming that our reaction conditions maintain a single-hit regime.

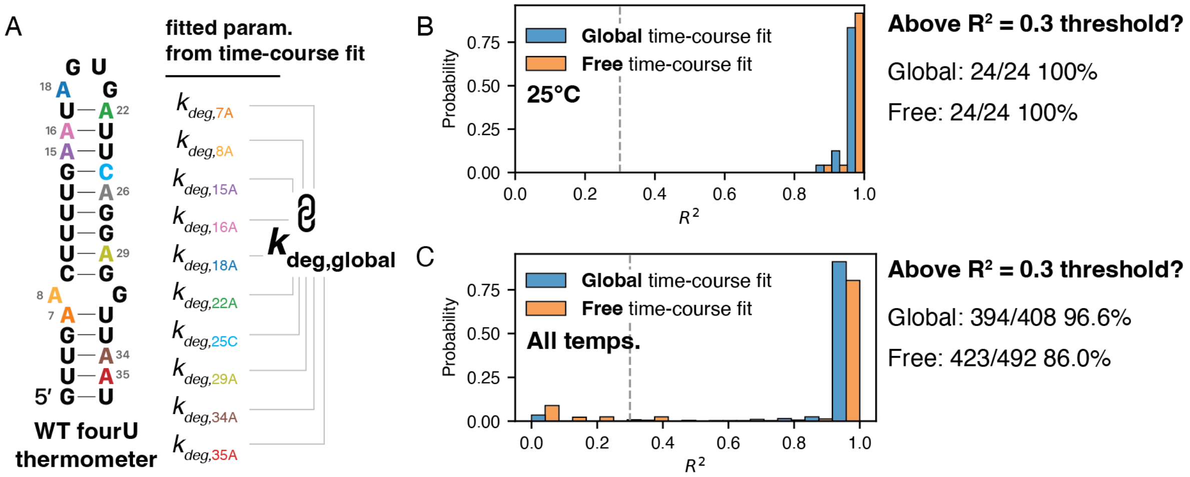


**Figure S4. Comparison of free vs. global time-course fit.** (**A**) Simultaneous fitting of time-courses across all sites yields a global $k_{\mathrm{deg}}$ estimate. This figure shows comparison of time-course fits for WT fourU RNA. (**B**) Effect of global fit on time-course goodness-of-fit ($R^2$) at 25°C, matching Fig. 1F, and (**C**) across all probed temperatures for the same construct. Annotated text show counts of data points that pass the quality control $R^2$ threshold of 0.3.

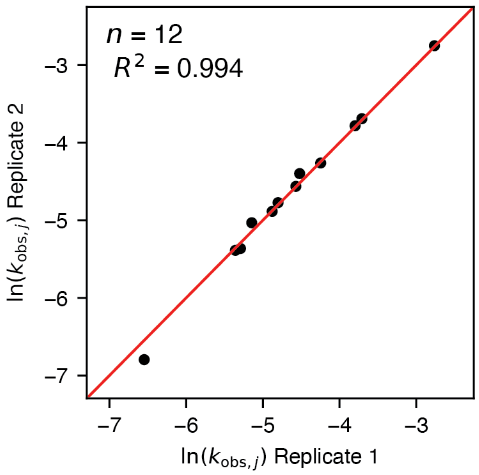


**Figure S5. Reproducibility of time-course chemical probing kinetics across independent replicates.** While Fig. 1F demonstrates that individual time-courses are well fit by the kinetic model, this figure compares parameter fits from independent experimental replicates for the same nucleotide sites, highlighting the reproducibility of the measurements.

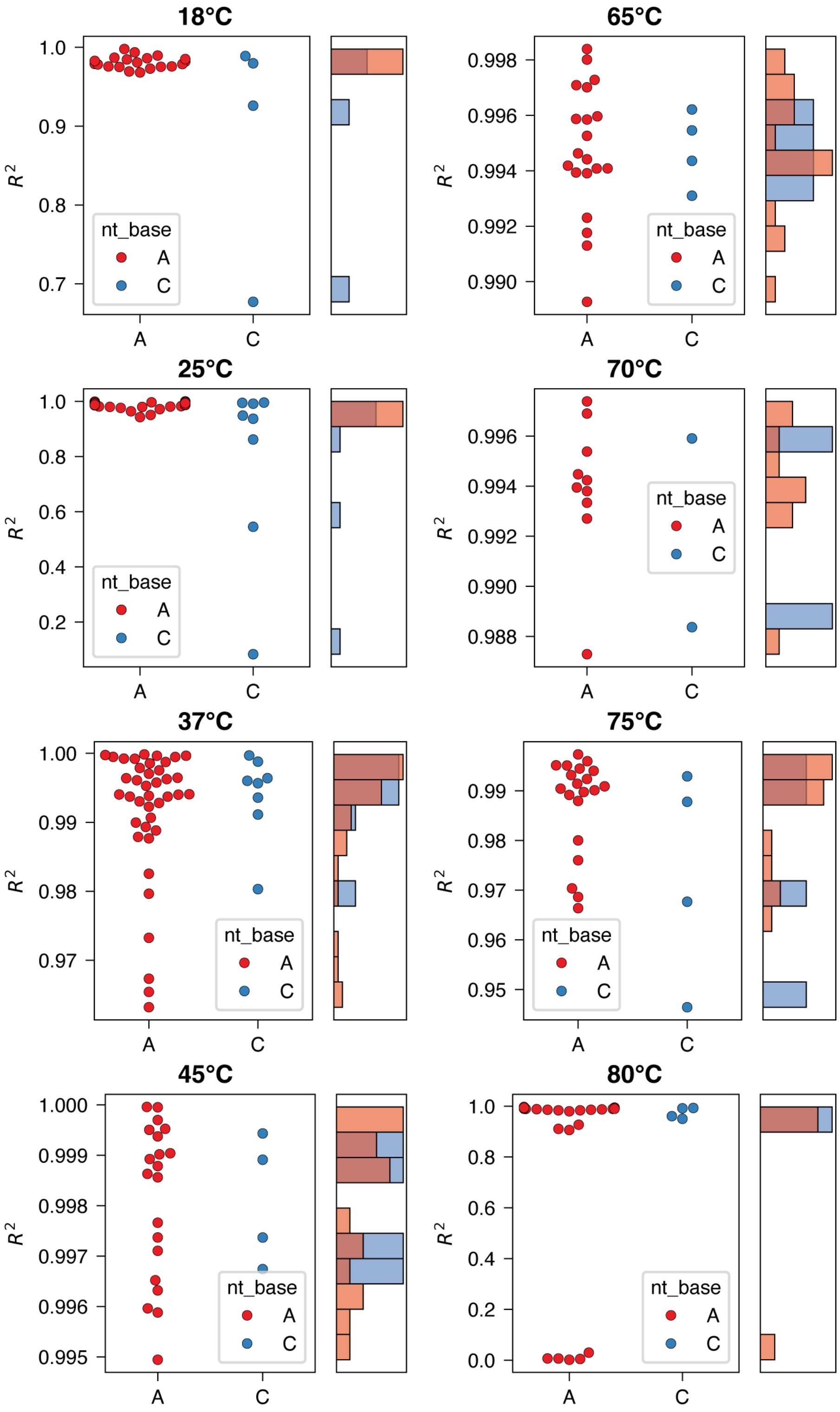


**Figure S6. Fit quality for time-course probing data of WT fourU thermometer across broad temperatures.** Swarm plots of $R^2$ from non-linear regression of time-course probing data (2-3 replicates) for A and C nucleotides. Histograms share a common y-axis with the swarm plots.

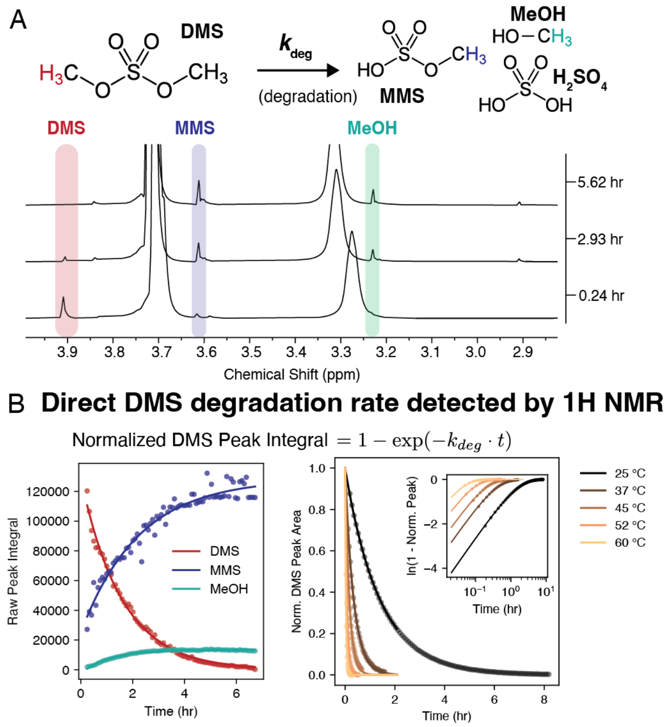


**Figure S7. Independent measurement of probe degradation reaction through time-resolved $^1$H NMR**. (**A**) DMS reacts with water to produce monomethyl sulfate (MMS) and methanol (MeOH), leading to an exponential decay of DMS concentration over time. Time-resolved $^1$H NMR tracks DMS degradation over time. (**B**) Quantification of NMR measurements show an exponential decay of DMS signal and increasing signal of MMS and MeOH. Measured decay at five different temperatures shows decay rate increases with temperature as expected. Inset shows $\ln(1 - \text{norm peak})$, supporting that these trajectories follow first-order exponential decay.

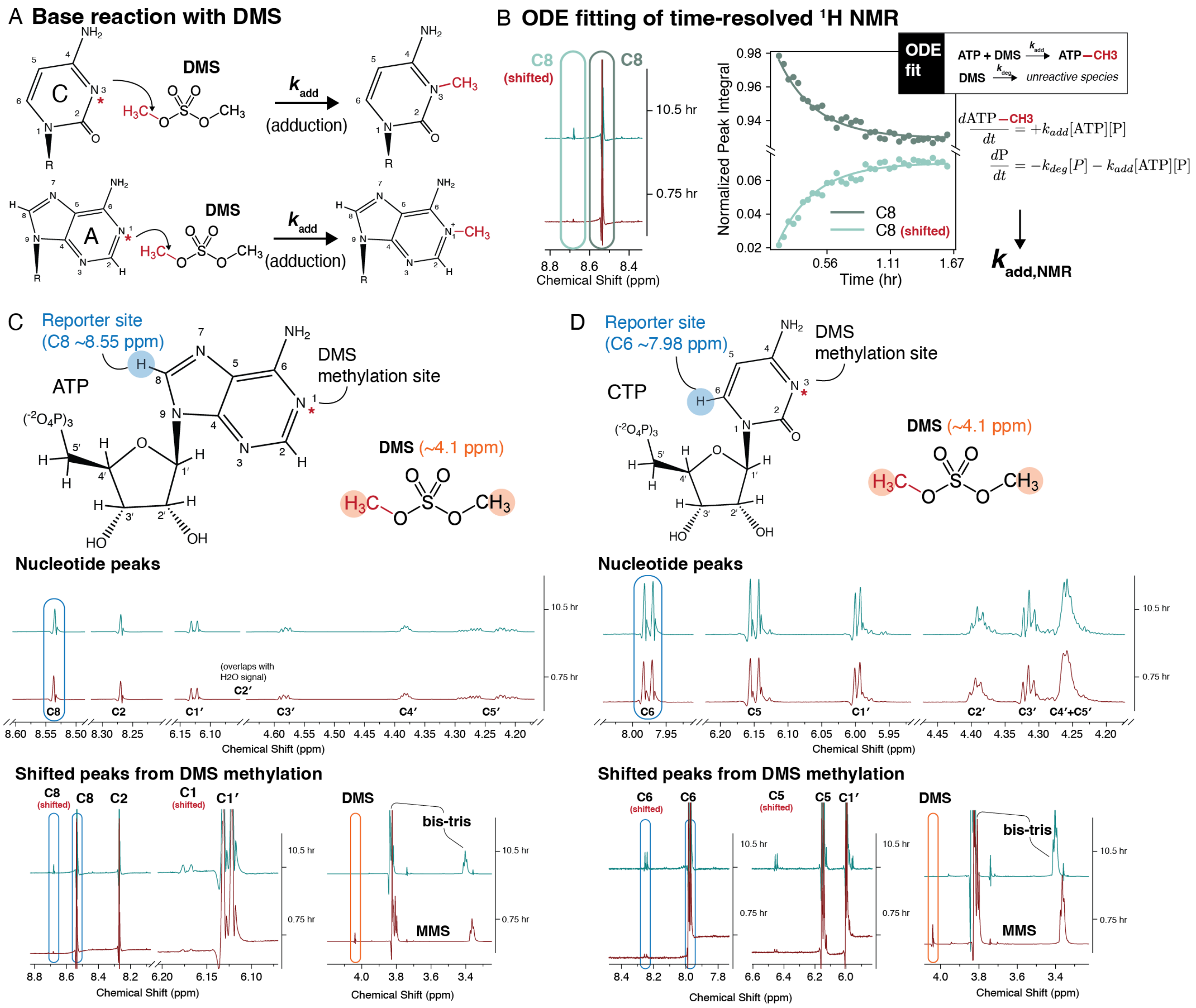


**Figure S8. Independent measurement of adduction formation reaction through time-resolved $^1$H NMR**. (**A**) Reaction schematic for DMS methylation of ATP and CTP at the N1 and N3 positions, respectively. (**B**) Representative time-resolved $^1$H NMR traces of ATP showing the C8 proton as a reporter of N1 methylation, indicated by the emergence of a downfield-shifted peak (labeled 'shifted' in red). Simultaneous fitting of ATP-$CH_3$ signal gain, ATP signal loss, and DMS ([P]) degradation using an ODE-based kinetic model yields the free-nucleotide adduction rate constant, $k_{\text{add}}$. (**C**) (top) ATP reaction products with DMS, annotated with observable protons in the $^1$H NMR spectrum at 0.75 hr and 10.5 hrs. Observable protons are annotated, and spectra were acquired in buffer conditions matched to fourU RNA probing experiments. (Bottom) Expanded view highlighting peaks that shift upon methylation. Reporter peaks used for kinetic fitting (C8 and C8-shifted) are highlighted in blue, and the DMS peak is highlighted in orange. (**D**) Corresponding $^1$H NMR spectra of CTP reaction products.

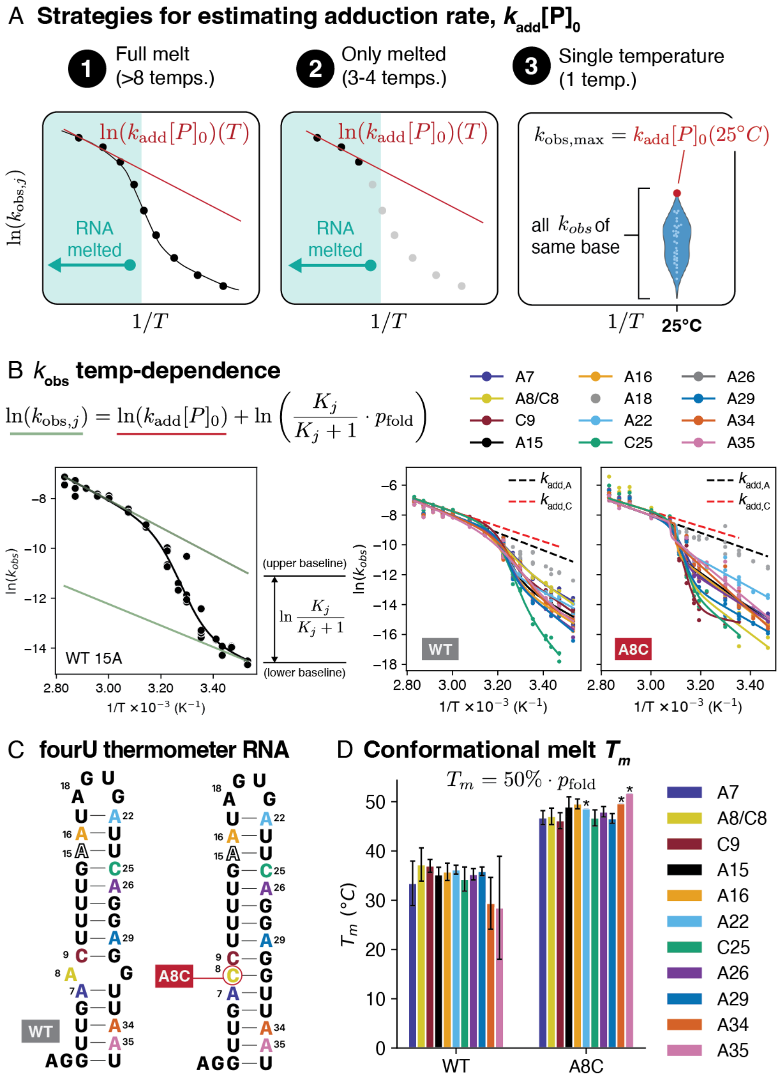


**Figure S9. DMS-derived probing kinetics exhibit two-state, temperature-dependent kinetics governed by RNA structural transitions.** (**A**) Strategies for estimating $k_{\text{add}}[P]_0$ from $k_{\text{obs,j}}$ as described in Supplementary Note 2. (**B**) Representative $\ln k_{\text{obs}}$ fit to a two-state model for position A15 of the WT fourU thermometer (Supplementary Note 2). Right: Two-state model fits for all A and C sites in WT and A8C fourU RNA. The aggregate $\ln(k_{\text{add}})$ curve extracted from fully melted data is shown as a dashed black (WT) or red (A8C) line and serves as a constrained upper baseline. (**C**) Secondary structures of the fourU htrA thermometer (WT and A8C variant), with A and C positions color-coded to match panels (B) and (D). (**D**) Conformational melting temperatures ($T_m$) extracted using Eq. 18 (Supplementary Note 2, Strategy 1) for selected A sites, grouped by WT and A8C constructs. Error bars represent standard errors from non-linear two-state fits. Asterisks (*) indicate sites for which fit uncertainties could not be estimated.

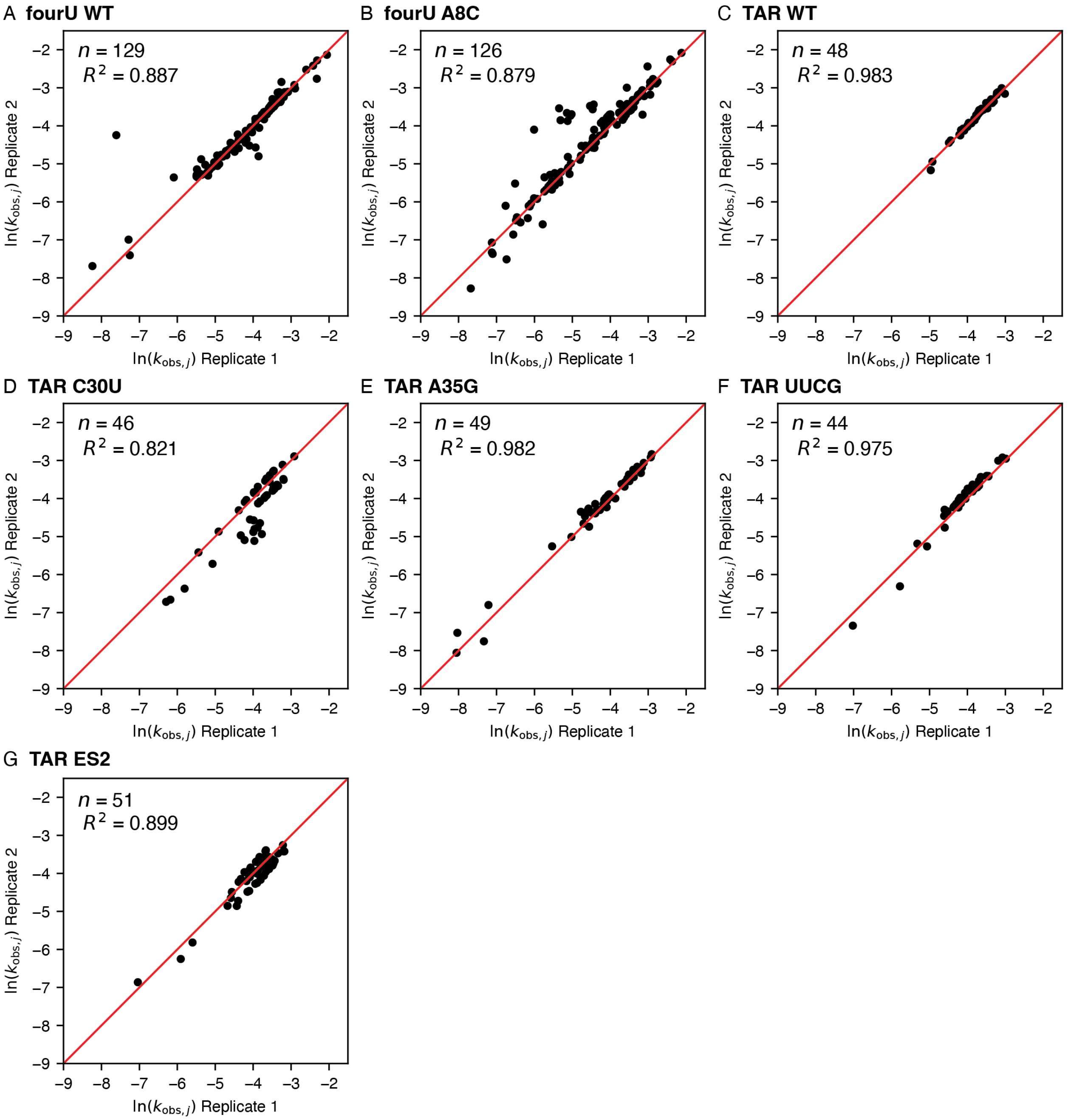


**Figure S10. Reproducibility of time-course fits across independent replicate time-course probing experiments of fourU and HIV-1 TAR RNAs.** Data points passing quality control ($R^2$ > 0.3) for fourU (**A-B**) and HIV (**C-G**) systems including reactions across all temperatures probed.

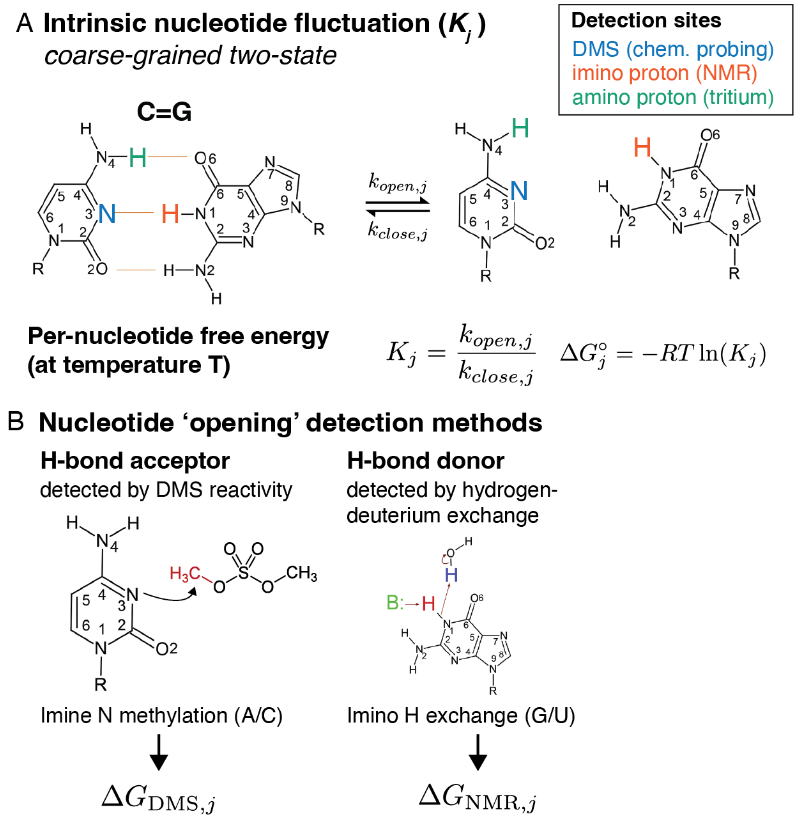


**Figure S11. Reference per-nt energies used to compare extracted $\Delta G^\circ_{\mathrm{DMS}}$.** (**A**) Nucleotide fluctuation is modeled as an equilibrium between two coarse-grained states: 'unreactive' (constrained) and 'reactive' (accessible). Different experimental methods probe distinct atomic sites within this framework. In the C=G canonical pair example shown, DMS and tritium-exchange report on the N3 and exocyclic amino positions of cytosine, respectively, whereas imino-proton exchange monitors the N1-H of guanine. (**B**) The extent of structural fluctuation required to reach the reactive configuration depends on the detection method. In DMS probing, the reactive state is probed by nucleophilic attack of DMS at the imine nitrogen (N1 in A, N3 in C (shown)) (*1*). In imino proton exchange detected by NMR, base-catalyzed (B:) deprotonation of the imino proton during transient base-pair opening enables exchange with solvent (*2*).

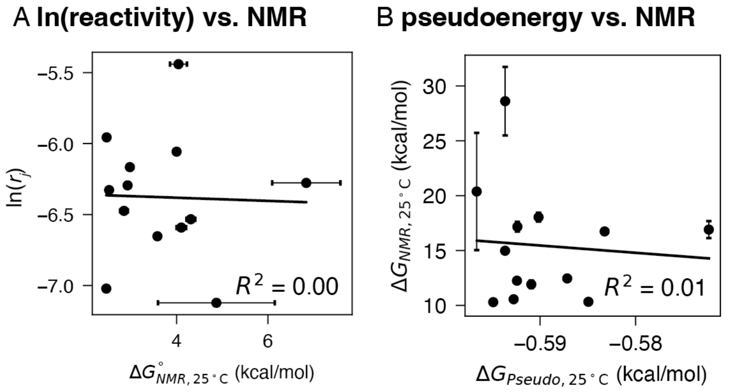


**Figure S12. Correlations between raw reactivities, pseudo-free energies, and $\Delta G_{NMR}$.** (**A**) Correlation between raw reactivities and NMR show no relationships. (**B**) In many studies, reactivities are converted into pseudoenergies with the heuristic $\Delta G^\circ_{pseudo,j} = -m\ln(r_j + 1) + b$ to constrain the prediction of secondary structure (*3*). Comparison of pseudoenergies (parametrized by default $m = 1.8$ kcal/mol and $b = -0.6$ kcal/mol taken from RNAstructure documentation) with $\Delta G^\circ_{\mathrm{NMR}}$ show no relationship.

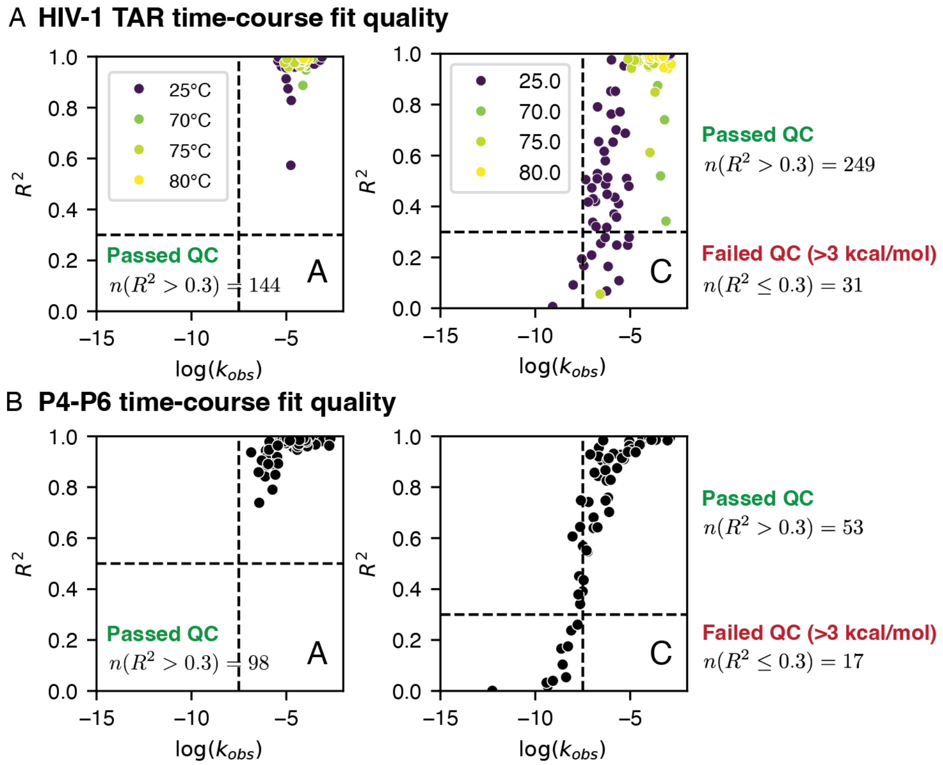


**Figure S13. Quality control of time-course fits for HIV-1 TAR and P4-P6 with respect to $\ln(k_{obs})$ values.** Scatter plots show R² values versus $\ln(k_{obs})$ for non-linear time-course fits at individual nucleotide sites. Panels correspond to HIV-1 TAR A sites (**A**), HIV-1 TAR C sites (**B**) across all mutant variants at 4 temperatures, P4-P6 A sites (**C**), and P4-P6 C sites (**D**) with and without $Mg^{2+}$. The horizontal black dashed line indicates the QC threshold ($R^2 = 0.3$). The number of data points in each panel is annotated. We observed that sites with poor fits ($R^2 < 0.3$) occurred when $\ln(k_{obs}) < -7.5$ (vertical black dashed line), which corresponded to positions having high stabilities ($\Delta G_{DMS} > 3$ kcal/mol). This threshold follows from $\ln(k_{obs}) = -7.5$ and $\ln(k_{add}[P]_0) = -2.7$, which together give $\ln\big(K/(K+1)\big) = -4.8$ corresponding to $\Delta G^{\circ}_{DMS} = 2.86$ kcal/mol.

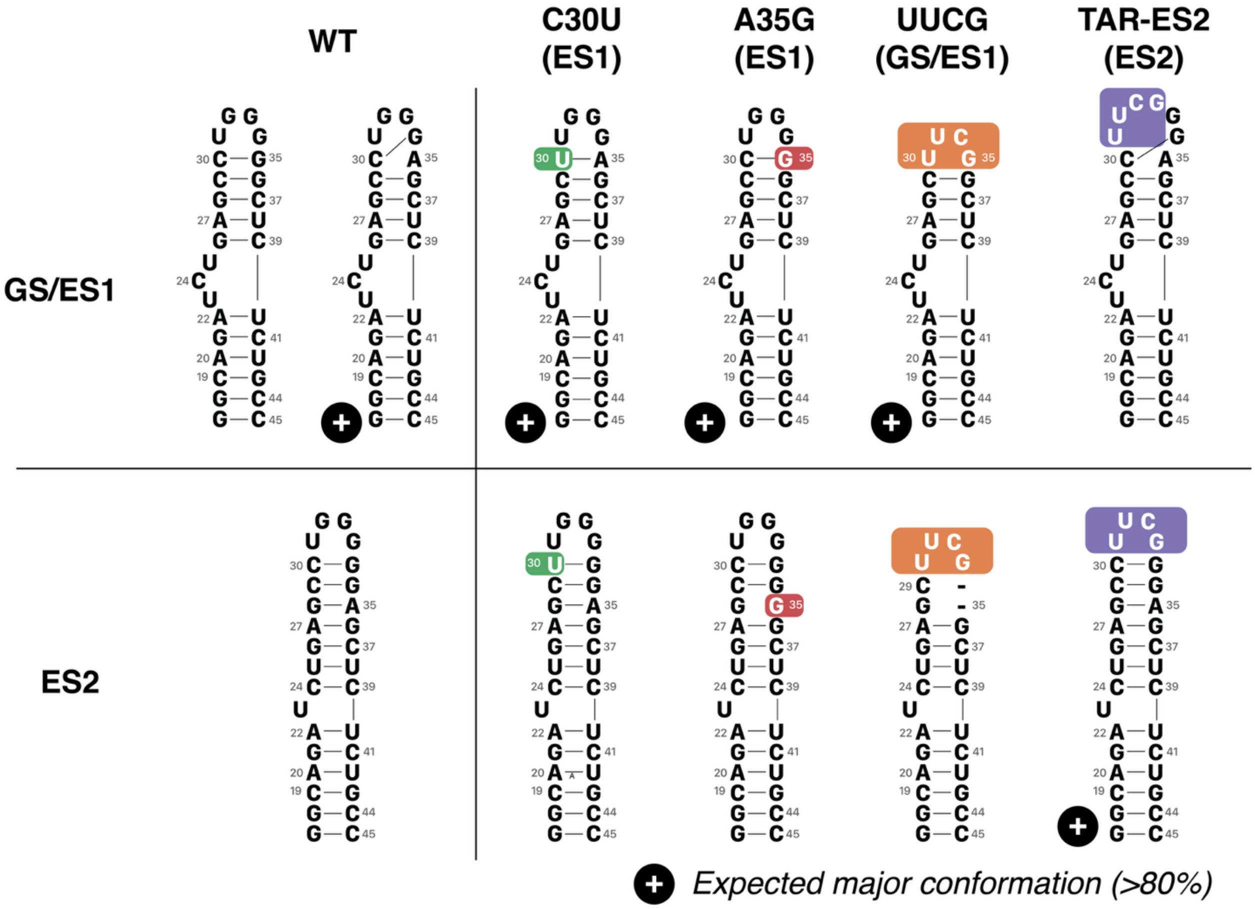


**Figure S14. Secondary structures of HIV-1 TAR mutants analyzed in this study.** Secondary structures are shown for each mutant mapped onto the GS/ES1 conformation (top) or the ES2 conformation (bottom). The expected dominant conformation for each mutant, as determined from prior NMR studies (*4-8*), is indicated.

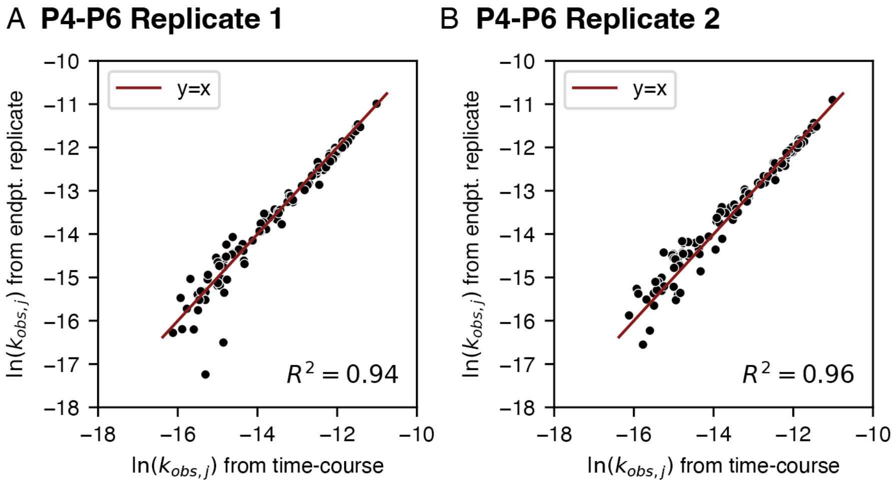


**Figure S15. Reproducibility of fitted $k_{obs,j}$ values for P4–P6 time-courses.** Correlation plots comparing $\ln(k_{obs,j})$ values obtained from time-course fits with values derived from two independent end-point replicates for the P4–P6 construct measured without $Mg^{2+}$.

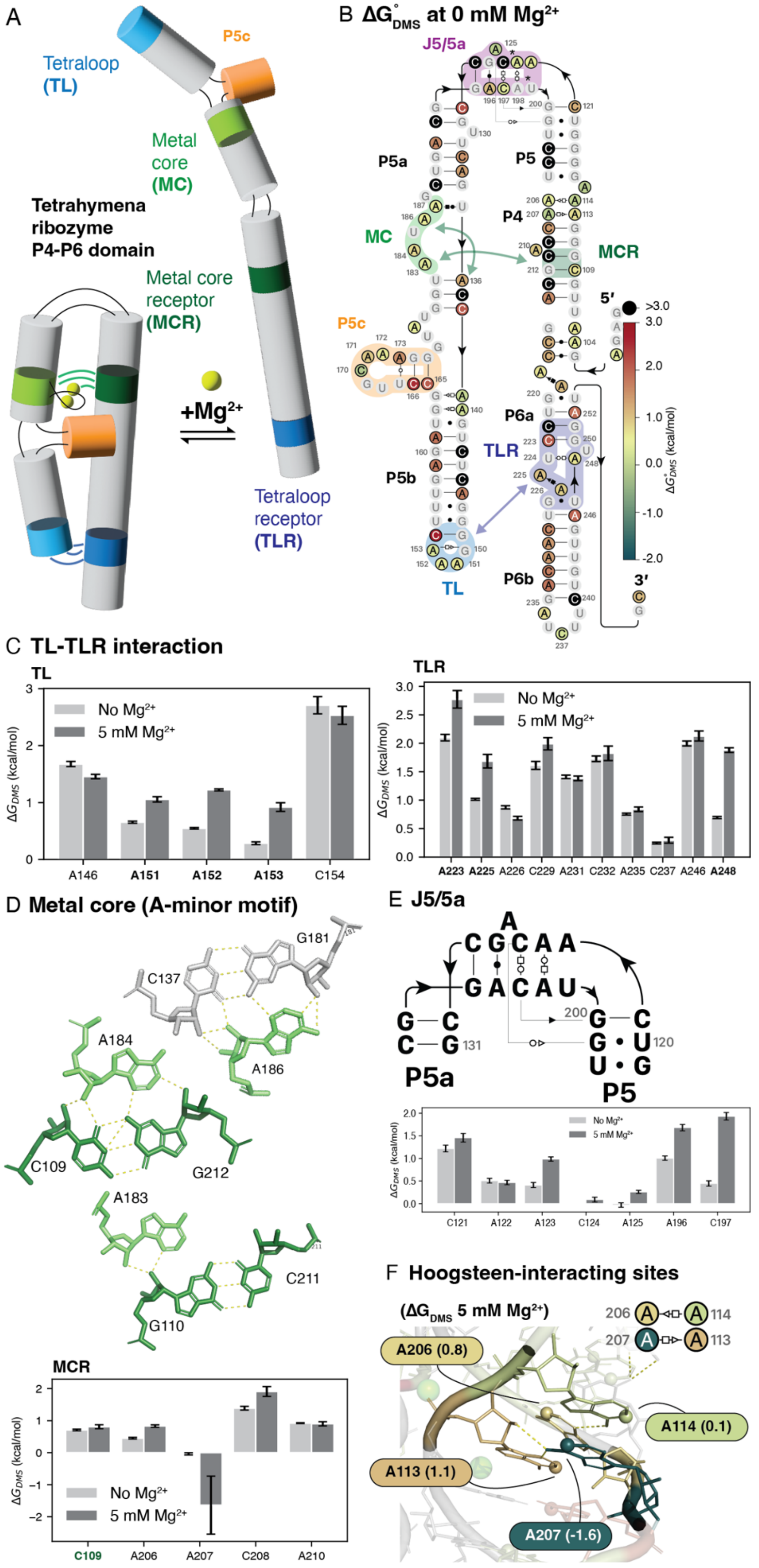
A
Tetraloop (TL)
P5c
Metal core (MC)
Tetrahymena ribozyme P4-P6 domain
Metal core receptor (MCR)
+Mg2+
Tetraloop receptor (TLR)
B ΔG°DMS at 0 mM Mg2+
J5/5a
P5a
P5
P4
MC
MCR
P5c
P6a
TLR
P5b
P6b
TL
5′
3′
>3.0
ΔG°DMS (kcal/mol)
C TL-TLR interaction
TL
TLR
No Mg2+
5 mM Mg2+
ΔGDMS (kcal/mol)
A146
A151
A152
A153
C154
A223
A225
A226
C229
A231
C232
A235
C237
A246
A248
D Metal core (A-minor motif)
G181
C137
A184
A186
C109
G212
A183
C211
G110
MCR
C109
A206
A207
C208
A210
E J5/5a
P5a
P5
C121
A122
A123
C124
A125
A196
C197
F Hoogsteen-interacting sites
(ΔGDMS 5 mM Mg2+)
A206 (0.8)
A114 (0.1)
A113 (1.1)
A207 (-1.6)

**Figure S16. P4-P6 system and $\Delta G^{\circ}_{\mathrm{DMS}}$ at specific contexts.** (**A**) Cartoon depiction of P4-P6 folding in the presence of $Mg^{2+}$. (**B**) Annotated secondary structure of P4-P6 with base pairs classified using Leontis-Westhof geometry. The tetraloop-tetraloop receptor (TL-TLR), J5/5a junction, P5c, and metal core-metal core receptor (MC-MCR) regions are highlighted. Nucleotides are colored according the $\Delta G^{\circ}_{\mathrm{DMS}}$ values at 0 mM Mg2+. (**C**) TL-TLR interaction network with corresponding $\Delta G^{\circ}_{\mathrm{DMS}}$ values for relevant nucleotides. (**D**) Metal core and A-minor interaction network with associated $\Delta G^{\circ}_{\mathrm{DMS}}$ values. All error bars represent standard error from non-linear fit of time-course profiles. (**E**) Secondary structure of the J5/5a junction annotated with non-canonical base pairs using Leontis-Westof classification with corresponding $\Delta G^{\circ}_{\mathrm{DMS}}$ bar plot. (**F**) Representative non-canonical Hoogsteen interaction sites annotated using Leontis-Westhof notation. Spheres are shown at the N1 position of adenines and are color-coded by $\Delta G^{\circ}_{\mathrm{Mg}}$ values.

### Supplementary Text

#### Supplementary Note 1. Derivation of a kinetic model for per-nucleotide chemical probing reactivities

Reaction overview

$$U_j \underset{k_{\text{close,j}}}{\overset{k_{\text{open,j}}}{\rightleftharpoons}} R_j$$

$$R_j + P \xrightarrow{k_{\text{add}}} M_j$$

$$P \xrightarrow{k_{\text{deg}}} \emptyset$$

In this model, we assume that each individual RNA nucleotide, $j$, fluctuates between a chemical probe-reactive (i.e. 'unstructured'), $R_j$, and a probe unreactive (i.e. 'structured'), $U_j$, state. The opening and closing rates of these fluctuations are given by $k_{\text{open,j}}$ and $k_{\text{close,j}}$. In an RNA structure probing experiment, the chemical probe, $P$, only reacts with a nucleotide in the reactive state to form a modified nucleotide, $M_j$, with an adduction rate constant, $k_{\text{add}}$. In parallel, the chemical probe, $P$, also hydrolyzes with water with degradation rate, $k_{\text{deg}}$. These reactions can be expressed as a system of ODEs:

$$\frac{dU_j}{dt} = -k_{\text{open}}[U_j] + k_{\text{close}}[R_j] \tag{1}$$

$$\frac{dR_j}{dt} = k_{\text{open}}[U_j] - k_{\text{close}}[R_j] - k_{\text{add}}[R_j][P] \tag{2}$$

$$\frac{dM_j}{dt} = k_{\text{add}}[R_j][P] \tag{3}$$

$$\frac{dP}{dt} = -k_{\text{deg}}[P] - k_{\text{add}}[R_j][P] \tag{4}$$

The resulting data from a chemical probing experiment are in the form of "reactivity", $r_j$, defined as the fraction of that nucleotide modified, or:

$$r_j = \frac{[M_j]}{[T_j]}, \tag{5}$$

where

$$[T_j] = [U_j] + [R_j] + [M_j], \tag{6}$$

is the total amount of RNA in the experiment.

Our goal is to use this model to derive an integrated rate equation for reactivity as a function of experimental- and probe-specific parameters ($[P]$, $[T]$, $k_{\text{add}}$, and $k_{\text{deg}}$) and the intrinsic RNA structural fluctuation parameters ($k_{\text{open}}$, $k_{\text{close}}$).

Model simplifications based on probing regimes

We make the following simplifications that are reasonable based on known RNA biophysics and the parameter regimes of typical probing experiments.

*Position independence.*

Given the concentration regime of our experiment, with probe concentration ([P] = 15.64 mM) being >> than the RNA concentration ([RNA] = 1 µM), RNAs are probed in a single-hit kinetic regime (Supplementary Figure S3). This allows Eqns. (1)-(4) to be used to model each position in the RNA independently. This means that $k_{\text{add}}$ is base-specific, but not RNA position specific. For simplicity, we drop the subscript $j$ in the rest of this derivation until the general result.

*Nucleotide fluctuation equilibrium assumption.*

The rates of nucleotide opening and closing are much faster than the rate of modification, $k_{\text{open}}, k_{\text{close}} \gg k_{\text{add}}$ such that nucleotide fluctuation equilibrium is maintained throughout the reaction. Here, a nucleotide "opening" and "closing" is a coarse-graining of all the motions that an RNA nucleotide experiences as it fluctuates between chemically reactive and unreactive states including base pair breathing, base stacking, backbone flexibility, helical fraying, tertiary contact engagement, and more. These motions are known to occur within the range of $10^{-12}$ to $10^{0}$s time-scales (*9-11*). The adduction reaction timescale, for DMS, is on the order of $10^{4}$s at 37°C. Under these assumptions, it follows that multiple opening events occur for every single modification event. Under this simplification, the intrinsic nucleotide fluctuations maintain equilibrium such that $\frac{dU}{dt} = \frac{dR}{dt} = 0$. Using these equations, we can derive a formula for the per-nucleotide fluctuation equilibrium constant, K:

$$K \equiv \frac{k_{\text{open}}}{k_{\text{close}}} = \frac{[R]}{[U]} \tag{7}$$

*Hydrolysis-mediated decay dominates chemical probe consumption.*

Under our experimental regime, $P$ is primarily consumed by hydrolysis due the large excess of reagent typically used in probing experiments. For example, in our experiments, 1 µM RNA is probed in >10 mM chemical probe and 55.6 M $H_2O$. Therefore, we make the simplification that $k_{\text{deg}}[P] \gg k_{\text{add}}[R][P]$. This allows us to integrate the first-order exponential decay rate ODE for $P$ (equation (4)) independently of the other equations to obtain a solution for $[P]$ as a function of reaction time:

$$\begin{aligned} \frac{dP}{dt} &= -k_{\text{deg}}[P] - k_{\text{add}}[R][P] \\ &\approx -k_{\text{deg}}[P] \end{aligned}$$

$$\int_{[\mathrm{P}]_0}^{[\mathrm{P}]} \frac{1}{[\mathrm{P}]} \cdot dP = \int_0^t -k_{\text{deg}} \cdot dt$$

$$\ln([\mathrm{P}]) - \ln([\mathrm{P}]_0) = -k_{\text{deg}}\, t$$

$$[P] = [P]_0 \cdot \exp(-k_{\text{deg}} \cdot t) \tag{8}$$

where $[P]_0$ is the amount of probe at the beginning of the reaction ($t = 0$).

Model solution

We seek a solution for

$$r(t) = \frac{[M]}{[T]}. \tag{9}$$

We start with Eq. (3)

$$\frac{dM}{dt} = k_{\text{add}}[R][P]$$

and find an expression for $[R]$ in terms of $[M]$. Using Eq. (6), and the definition of $K$ (Eq. (7)), we substitute $[R] = [T] - [M] - [U]$ to derive an expression for $[U]$ and $[R]$.

$$\begin{aligned} K &= \frac{[T] - [M] - [U]}{[U]} \\ [U] &= \frac{1}{K}([T] - [M] - [U]) \\ [U] + \frac{[U]}{K} &= \frac{1}{K}([T] - [M]) \\ [U]\left(1 + \frac{1}{K}\right) &= \frac{1}{K}([T] - [M]) \\ [U]\left(\frac{K+1}{K}\right) &= \frac{1}{K}([T] - [M]) \\ [U] &= \frac{1}{K+1}([T] - [M]) \end{aligned}$$

We then derive an equation for $[R]$:

$$\begin{aligned} [R] &= [U]K \\ &= \left(\frac{1}{K+1}([T] - [M])\right)K \\ &= \frac{K}{K+1}([T] - [M]). \end{aligned}$$

Using expressions for $[R]$ and $[P]$ (Eq. 8), we can now derive an expression for $\frac{dM}{dt}$:

$$
\begin{aligned}
\frac{dM}{dt} &= k_{\text{add}}[R][P] \\
&= k_{\text{add}}\left(\frac{K}{K+1}([T]-[M])\right)\left([P]_0 \cdot \exp\left(-k_{\text{deg}} \cdot t\right)\right) \\
&= ([T]-[M])\left(\frac{K}{K+1}\right) k_{\text{add}}[\text{P}]_0 \cdot \exp\left(-k_{\text{deg}} \cdot t\right)
\end{aligned}
$$

Separating variables, we obtain:

$$
\left(\frac{1}{([T]-[M])}\right) \cdot dM = \left(\frac{K}{K+1}\right) k_{\text{add}}[\text{P}]_0 \cdot \exp\left(-k_{\text{deg}} \cdot t\right) \cdot dt \tag{10}
$$

Integrating both sides, we get:

$$
\int_{[M]_0}^{[M]} \left(\frac{1}{([T]-[M])}\right) \cdot dM = \int_0^t \left(\frac{K}{K+1}\right) k_{\text{add}}[P]_0 \cdot \exp\left(-k_{\text{deg}} \cdot t\right) \cdot dt \tag{11}
$$

We now solve each side separately. Using $[M]_0 = 0$, we have:

$$
\begin{aligned}
\text{LHS} &= \int_{[M]_0}^{[M]} \left(\frac{1}{([T]-[M])}\right) \cdot dM \\
&= -(\ln([T]-[M]) - \ln([T])) \\
&= -\ln\left(\frac{[T]-[M]}{[T]}\right)
\end{aligned}
$$

$$
\begin{aligned}
\text{RHS} &= \int_0^t \left(\frac{K}{K+1}\right) k_{\text{add}}[\text{P}]_0 \cdot \exp\left(-k_{\text{deg}} \cdot t\right) \cdot dt \\
&= -\frac{K}{K+1}\frac{k_{\text{add}}}{k_{\text{deg}}}[\text{P}]_0 \cdot \left(\exp\left(-k_{\text{deg}} \cdot t\right) - \exp(0)\right) \\
&= \frac{K}{K+1}\frac{k_{\text{add}}}{k_{\text{deg}}}[\text{P}]_0 \cdot \left(1 - \exp\left(-k_{\text{deg}} \cdot t\right)\right)
\end{aligned}
$$

Combining both sides, and using the definition of $r(t)$ (Eq. 9), we have:

$$\text{LHS} = \text{RHS}$$

$$-\ln\left(\frac{[T]-[M]}{[T]}\right) = \frac{K}{K+1}\frac{k_{\text{add}}}{k_{\text{deg}}}[P]_0 \cdot \left(1 - exp\left(-k_{\text{deg}} \cdot t\right)\right)$$

$$\frac{[T]-[M]}{[T]} = \exp\left(-\frac{K}{K+1}\frac{k_{\text{add}}}{k_{\text{deg}}}[P]_0 \cdot \left(1 - exp\left(-k_{\text{deg}} \cdot t\right)\right)\right)$$

$$[M] = [T] - [T]exp\left(-\frac{K}{K+1}\frac{k_{\text{add}}}{k_{\text{deg}}}[P]_0 \cdot \left(1 - exp\left(-k_{\text{deg}} \cdot t\right)\right)\right)$$

$$[M] = [T]\left(1 - \exp\left(-\frac{K}{K+1}\frac{k_{\text{add}}}{\text{k}_{\text{deg}}}[P]_0 \cdot \left(1 - exp\left(-k_{\text{deg}} \cdot t\right)\right)\right)\right)$$

Dividing both sides by $[T]$:

$$r(t) = \frac{[M]}{[T]} = 1 - \exp\left(-\frac{K}{K+1}\frac{k_{\text{add}}}{k_{\text{deg}}}[P]_0 \cdot \left(1 - \exp\left(-k_{\text{deg}} \cdot t\right)\right)\right) \tag{12}$$

For each nucleotide at site $j$, reactivity as a function of time is given by:

$$r_j(t) = \frac{[M_j]}{[T_j]} = 1 - \exp\left(-\frac{K_j}{K_j+1}\frac{k_{\text{add}}}{k_{\text{deg}}}[P]_0 \cdot \left(1 - \exp\left(-k_{\text{deg}} \cdot t\right)\right)\right). \tag{13}$$

In this model, reactivity is a complex function of experimental parameters ($[P]_0$, $k_{\text{add}}$, $k_{\text{deg}}$, $t$) and intrinsic information about RNA structure ($K_j$), where the kinetic parameters and equilibrium constant all have distinct temperature-dependences.

This equation is used to fit time-course reactivity profiles (Methods), which yields 2 fit parameters, $k_{\text{deg}}$ and $k_{\text{obs},j}$:

$$k_{\text{obs},j} = \frac{K_j}{K_j+1}\, k_{\text{add}}[P]_0 \tag{14}$$

*Code implementation*

The full implementation of this model is available in the *nerd* GitHub repository (see `pipeline/plugins/timecourse/baseline.py`, function `_fmod_model`). This implementation extends the original formulation by explicitly incorporating a baseline reactivity term, $r_{j,0}$, which accounts for background signal at $t = 0$. Under this formulation, the time-dependent fraction of modified nucleotides is given by:

$$r_j(t) = \frac{[M_j]}{[T_j]} = 1 - \exp\left(-\frac{K_j}{K_j + 1}\frac{k_{\text{add}}}{k_{\text{deg}}}[P]_0 \cdot (1 - \exp(-k_{\text{deg}} \cdot t))\right) + r_{j,0}.$$

**Supplementary Note 2. Extracting $K_j$ from $k_{\text{obs}}$ by estimating adduct formation rate, $k_{\text{add}}$.**

In a time-course experiment, $k_{\text{obs, j}}$ values are fitted for each nucleotide. As seen from Eq. 14, $k_{\text{obs, j}}$ contains a contribution from the adduct formation rate, $k_{\text{add}}[P]_0$, and intrinsic information about RNA structure, $K_j$, each of which depend on temperature. Our goal is to establish a method to separate these two contributions to extract $K_j$ at specific temperatures.

Following Eq. 14, we have:

$$\ln\left(\frac{K_j(T)}{K_j(T) + 1}\right) = \ln\left(k_{\text{obs,j}}(T)\right) - \ln(k_{\text{add}}(T)[P]_0), \tag{15}$$

where we have added explicit temperature dependence. Our first step therefore is to estimate the adduction rate constant, $k_{\text{add}}(T)$ to be able to perform the required subtraction, noting that this must be done for each temperature studied. We can do this by examining regimes in which nucleotide positions are fully unfolded. In this regime, these nucleotides are in the fully reactive state, such that $K_j \to \infty$ and $\frac{K_j}{K_j+1} \to 1$, which according to Eq. 14 $k_{\text{obs,j}} \to k_{\text{add}}[P]_0$. Therefore, the adduction rate constant can be obtained directly from $k_{\text{obs,j}}$ when nucleotides are fully unfolded.

There are two experimental scenarios in which nucleotides can be assumed to be fully unfolded: (i) when a complete melting curve is measured, and (ii) when data are collected only at temperatures above the melting transition. These regimes motivate the three strategies used in this study to estimate $k_{\text{add}}(T)$ under different experimental constraints, including the number of temperatures sampled, the melting behavior of the RNA, and the overall complexity of the system, as described below (fig. S9A).

<u>Strategy 1: Using full two-state melting curve with measurements over 8-10 temperatures.</u>

For simple RNAs exhibiting a cooperative two-state thermal transition (e.g. fourU hairpin), we can obtain a complete melting curve across 8-10 temperatures (fig. S9).

The upper-temperature baseline of the melt corresponds to the regime where structural constraints vanish, and $k_{\text{obs,j}} \to k_{\text{add}}[P]_0$. Here

$$\ln(k_{\text{obs,melted}}) = \ln(k_{\text{add}}(T)) + \ln([P]_0).$$

If $k_{\text{add}}(T)$ follows Arrhenius behavior across temperatures as expected, then:

$$k_{\text{add}}(\text{T}) = A \cdot \exp(-E_a/RT)$$

$$\ln(k_{\text{obs,melted}}) = -\frac{E_a}{R} \cdot \frac{1}{T} + \ln(A) + \ln([P]_0) \tag{16}$$

where $E_a$ is the activation energy and $A$ is the pre-exponential factor for $k_{\text{add}}$. Fig. 2D confirms linear Arrhenius behavior shown in $\ln(k_{\text{add}})$ vs. 1/T plots where concentration has been factored out.

The lower baseline corresponds to the RNA when it is fully folded. In this case, $K_j \ll 1$, and $k_{\text{obs,j}} \to K_j k_{\text{add}}[P]_0$ , and

$$\ln(k_{\text{obs,folded}}) = \ln(\text{K(T)}) + \ln(k_{\text{add}}(T)) + \ln([P]_0).$$

Using the relationship $\Delta G = \Delta H - T\Delta S = -\text{RT} \ln K$, we model the temperature dependence of $\ln(K)$ as:

$$\ln(K) = -\frac{\Delta H}{R} \cdot \frac{1}{T} + \frac{\Delta S}{R}.$$

We therefore have:

$$\ln(k_{\text{obs,folded}}) = -\left(\frac{\Delta H}{R} + \frac{E_a}{R}\right)\left(\frac{1}{T}\right) + \left(\frac{\Delta S}{R} + \ln(A)\right) + \ln([P]_0) \tag{17}$$

We can then model the complete curve of $\ln(k_{\text{obs}})$ vs $T$ by interpolating between these two baselines. We model the RNA as occupying 2 global conformational states (e.g. helix vs. random coil) with equilibrium constant (*12*):

$$K_{\text{conf}}(\text{T}) = \exp\left(\frac{\Delta H_{\text{conf}}}{R}\left(\frac{1}{T_M} - \frac{1}{T}\right)\right)$$

The fractional populations of folded and melted states are given by:

$$p_{\text{fold}}(T) = \frac{K_{\text{conf}}(T)}{K_{\text{conf}}(T) + 1}$$

$$p_{\text{melt}}(T) = 1 - p_{\text{fold}}(T) = \frac{1}{K_{\text{conf}}(T) + 1} \tag{18}$$

The observed rate constant $k_{\text{obs}}$ is then a weighted sum by fractional populations of each state:

$$\ln(k_{\text{obs}}(T)) = \ln(k_{\text{obs, melted}}(T)) \cdot p_{\text{melt}}(T) + \ln(k_{\text{obs, folded}}(T)) \cdot p_{\text{fold}}(T) \tag{19}$$

This results in a classic 2-state melting curve in $\ln(k_{\text{obs}})$. We use Eqns. 16, 17 and 18 together in Eq. 19 to simultaneously fit the combined parameters:

| **Parameter Name as Implemented** | **Variable** | **Description** | **Equation** |
|---|---|---|---|
| a | $\frac{E_a}{R}$ | Activation energy of adduction reaction, slope of upper baseline | Eq. 16 |

| | | | |
|---|---|---|---|
| b | $ln(A) + ln([P]_0)$ | Pre-exponential baseline of adduction reaction, intercept of upper baseline | Eq. 16 |
| c | $\frac{\Delta H}{R} + \frac{E_a}{R}$ | Local enthalpy of intrinsic nucleotide opening, slope of lower baseline | Eq. 17 |
| d | $\frac{\Delta S}{R} + \ln(A) + ln([P]_0)$ | Local entropy of intrinsic nucleotide opening, intercept of lower baseline | Eq. 17 |
| f | $\Delta H_{\text{conf}}$ | Conformational enthalpy, steepness or cooperativity of melting curve | Eq. 18 |
| g | $T_M$ | Conformational melting point | Eq. 18 |

These parameters allow us to extract $K$ using $\Delta H$ and $\Delta S$. More intuitively, subtracting the lower baseline from the upper baseline gives a protection factor, $K/(K+1)$ (fig. S9).

We use this approach when analyzing the fourU system.

Note: This method requires the RNA to behave as a two-state system. If additional intermediates or non-cooperative behavior are present, the model may be invalid.

*Code implementation*
The full implementation of this model is available in the *nerd* GitHub repository (see `pipeline/plugins/tempgrad/two_state.py`, function `melt_fit`).

Strategy 2: Using a linear fit at 3-4 temperatures where the RNA is fully melted.

A reduced but still robust strategy is to collect $\ln k_{\text{obs}}$ at 3-4 temperatures where the RNA is known to be fully melted (figure S9E, strategy 2). In this regime,

$$k_{\text{obs}}(T) \approx k_{\text{add}}(T)[P]_0.$$

If $k_{\text{add}}(T)$ follows Arrhenius behavior across temperatures as expected, then:

$$k_{\text{add}} = A \cdot e^{-E_a/RT}$$
$$\ln(k_{\text{add}}) = -\frac{E_a}{R} \cdot \frac{1}{T} + \ln(A) + \ln([P]_0)$$

We can therefore use the 3-4 temperatures, where RNA is melted, to fit $E_a$ and $A$. This gives the temperature-dependence of $k_{\text{add}}(T)$, which can be extrapolated to lower temperatures to extract $K/(K+1)$, using Eq. (15):

$$\ln\left(\frac{K_j(T)}{K_j(T)+1}\right) = \ln\left(k_{\text{obs},j}(T)\right) + \frac{E_a}{R}\cdot\frac{1}{T} - \ln(A) - \ln([P]_0).$$

Note: This approach assumes that the RNA melts completely at experimentally accessible temperatures and that structural contributions are negligible in this window.

We use this approach when analyzing the HIV-1 TAR system.

*Code implementation*

The full implementation of this model is available in the *nerd* GitHub repository (see `pipeline/plugins/tempgrad/arrhenius.py`, function `_linear_model`).

Strategy 3: Using a single-temperature estimate.

For large or highly stable RNAs that do not melt under experimentally feasible temperatures, direct observation of a melted upper baseline is not possible. In these cases, we approximate $k_{\text{add}}[P]_0$ using the maximum measured $k_{\text{obs,max}}$ value across experimental conditions, assuming that the most reactive nucleotides are effectively unstructured and thus closest to the intrinsic adduction limit. In this case,

$$\ln\left(\frac{K_j(T)}{K_j(T)+1}\right) = \ln\left(k_{\text{obs,j}}(T)\right) - \ln\left(k_{\text{obs,max}}(T)\right).$$

While this represents the least constrained estimate, it is often the only practical approach for complex RNAs whose folds remain stable across the full temperature range accessible to DMS probing. We use this approach for analyzing the P4-P6 system.

### Supplementary Note 3. Calculating $\boldsymbol{\Delta G^\circ_{\text{DMS}}}$ from $\boldsymbol{K_j}$.

To calculate $\Delta G^\circ_{\text{DMS}}$, we first isolate the equilibrium constant $K_j$ from the measured fractional open population, then apply the standard thermodynamic relation $\Delta G = -RT\ln K$. Applying this to each nucleotide gives:

$$\Delta G^\circ_{\text{DMS},j} = -RT\ln K_j.$$

Throughout this work, we use the degree symbol (°) to denote a standard-state free energy change ($\boldsymbol{\Delta G}$°). Here, the standard state is defined at the temperature of interest rather than a fixed reference temperature.

### Supplementary Note 4. Uncertainty sources and error propagation

Error propagation for extracting $\Delta G^\circ_{\text{DMS},j}$

We propagate uncertainties using first-order (Taylor) error propagation. For a scalar transformation $y = g(x)$ with small uncertainty $\sigma_x$,

$$\sigma_y \approx \left|\frac{dg}{dx}\right|\sigma_x,$$

and for independent uncertainties entering additively, variances add. Let

$$f_{R,j}(T) = \frac{K_j(T)}{K_j(T)+1}$$

be the fraction of the population of nucleotide $j$ is in the reactive state at temperature $T$.

From Eq. (15),

$$\ln f_{R,j}(T) = \ln k_{\mathrm{obs},j}(T) - \ln(k_{\mathrm{add}}(T)[P]_0).$$

We denote the fitted uncertainty in $\ln k_{\mathrm{obs},j}(T)$ as $\sigma_{1,j}$, and the uncertainty in $\ln(k_{\mathrm{add}}(T)[P]_0)$ as $\sigma_2$. Assuming these are independent,

$$\sigma_{\ln f_{R,j}} = \sqrt{\sigma_{1,j}^2 + \sigma_2^2}.$$

We next propagate $\sigma_{\ln f_{R,j}}$ to $f_{R,j}$. Let $x = \ln f$ and $f = e^x$. Then

$$\frac{df}{dx} = e^x = f \quad \Rightarrow \quad \sigma_f = \left|\frac{df}{dx}\right| \sigma_x = f\,\sigma_{\ln f}.$$

Thus,

$$\sigma_{f_{R,j}} = f_{R,j}\,\sigma_{\ln f_{R,j}}.$$

Using the definition of $f_{R,j}$, we solve for $K_j$:

$$K_j = \frac{f_{R,j}}{1 - f_{R,j}}.$$

Differentiating $K(f) = \frac{f}{1-f}$ gives

$$\frac{dK}{df} = \frac{1}{(1-f)^2} \quad \Rightarrow \quad \sigma_{K_j} = \left|\frac{dK}{df}\right| \sigma_{f_{R,j}} = \frac{\sigma_{f_{R,j}}}{\left(1 - f_{R,j}\right)^2}.$$

Finally, for $\ln K_j$,

$$\frac{d\ln K}{dK} = \frac{1}{K} \quad \Rightarrow \quad \sigma_{\ln K_j} = \frac{\sigma_{K_j}}{K_j}.$$

Substituting $K_j = \frac{f}{1-f}$ and $\sigma_f = f\sigma_{\ln f}$ yields a compact expression:

$$\begin{aligned}\sigma_{\ln K_j} &= \frac{1}{K_j} \cdot \frac{\sigma_{f_{R,j}}}{\left(1 - f_{R,j}\right)^2} \\ &= \frac{1}{\frac{f}{1-f}} \cdot \frac{f\sigma_{\ln f}}{(1-f)^2} \\ &= \frac{\sigma_{\ln f_{R,j}}}{1 - f_{R,j}}.\end{aligned}$$

Therefore, the uncertainty in the DMS-derived free energy $\Delta G^\circ_{\mathrm{DMS},j} = -RT \ln K_j$ is

$$\sigma_{\Delta G_j} = RT\,\sigma_{\ln K_j}$$

$$= RT\,\frac{\sigma_{\ln f_{R,j}}}{1 - f_{R,j}}$$

$$= RT\,\frac{\sqrt{\sigma_{1,j}^2 + \sigma_2^2}}{1 - f_{R,j}}.$$

*Code implementation*

The implementation of this is used in various notebooks in the *Choi_PRIME_chemprobing_2026* GitHub repository.

Error propagation between independent replicates

When averaging n independent replicate measurements with propagated uncertainties $\boldsymbol{\sigma_r}$, the uncertainty on the mean is computed by propagating errors through the arithmetic mean. For

$$\boldsymbol{\bar{x} = \frac{1}{n}\sum_{r=1}^{n} x_r,}$$

first-order error propagation for independent uncertainties gives

$$\boldsymbol{\sigma_{\bar{x}}^2 = \sum_{r=1}^{n}\left(\frac{\partial \bar{x}}{\partial x_r}\right)^2 \sigma_r^2}$$

$$\boldsymbol{= \sum_{r=1}^{n}\left(\frac{1}{n}\right)^2 \sigma_r^2}$$

$$\boldsymbol{= \frac{1}{n^2}\sum_{r=1}^{n} \sigma_r^2.}$$

Thus, the uncertainty of the mean is

$$\boldsymbol{\sigma_{\bar{x}} = \frac{\sqrt{\sum_{r=1}^{n} \sigma_r^2}}{n}.}$$

*Code implementation*

The implementation of this is used in various notebooks in the *Choi_PRIME_chemprobing_2026* GitHub repository.

**Supplementary Note 5. Design of RNA flanking sequence for mutational profiling and sequencing**

Modification detection by reverse transcription overview

Chemical probing detects RNA modifications through reverse transcription (RT), which requires a primer-binding site at the 3′-end of the RNA. Because RT priming occludes the primer-binding region and the immediately adjacent nucleotides, a spacer sequence is routinely included between the 3′ end of the target RNA and the RT primer site to ensure complete coverage of the target. Additional sequence is similarly appended to the 5′-end to prevent loss of information during downstream amplification. In RT-MaP workflows, PCR primers bind directly to full-length cDNA, and corresponding primer-binding sites are therefore incorporated at the RNA transcript level.

As the RNA is made and folded in the context of these flanking regions, it is important to ensure that the flanking sequences do not disrupt the target RNA folding. Because adjacent hairpins can have a stabilizing effect on RNA structures (*13*), we opted to design unstructured regions rather than flanking hairpin cassettes.

Designing unstructured flanking sequences

For the fourU thermometer and HIV-1 TAR RNAs, we used NUPACK (*14*) to automate a design pipeline for flanking sequences to minimally disrupt the target RNA. We implemented this pipeline using NUPACK 4.0.1.1 Python module on a Jupyter notebook (see **Data, code, and materials availability).** Briefly, this pipeline generates candidate sequences for both 5′- and 3′-ends across multiple temperatures, assesses the overall ensemble defect score of the designs (*14*), which represents a distance between the target design structure and the ensemble of folds a candidate sequence is predicted to fold into, evaluates PCR $T_m$ compatibility, and chooses an optimal design.

At each of 5 °C, 25 °C, and 37 °C we generated sequences that preserved the target RNA secondary structure, excluding predefined homopolymeric nucleotide motifs to avoid low-complexity sequences. For each design we evaluated the predicted ensemble defect and minimum free energy (MFE) fold, then extracted the first 18 nucleotides as the 5′ PCR primer and the final 24 nucleotides as the RT primer region. For HIV, using the IDT OligoAnalyzer API we further computed the reverse-complement and RNA melting temperature of the RT primer (trimming the complement by five nucleotides to set primer length) and queried the NEB thermodynamic API to obtain DNA melting temperatures for both primers. Candidates were retained only if their ensemble defect was below 0.02, the 5′/3′ primer melting temperatures differed by ≤5 °C, and the RT primer melting point exceeded 47 °C. The remaining sequences were reanalyzed with single-strand NUPACK ensemble calculations and sorted by ensemble defect. A winning candidate was chosen with the highest RT primer melting temperature with lowest ensemble defect.

For the P4-P6 construct, we based our flanking sequence design on a previous construct from (*15*), using similar primer sequences. We made one modification to add an additional 6-nt spacer the 5′-end, which we confirmed did not disrupt the target P4-P6 RNA fold using NUPACK.

*Code implementation*

Jupyter notebooks used for NUPACK design pipeline of fourU thermometer and HIV-1 TAR is available in the *Choi_PRIME_Chemprobing_2025* GitHub repository (see `Figure_analysis/Methods_construct_design/fourU_design.ipynb` and `Figure_analysis/Methods_construct_design/HIV_design.ipynb`).

**Supplementary Note 6. Preparation of temperature-corrected folding buffers**

Overview

Temperature changes shift buffer $pK_a$ values, such that a buffer prepared at room temperature, $pH_{\text{prep}}$, will not have the same $pH$ when used at a higher temperature. To maintain the same $pH$ across a range of temperatures, a temperature-dependence correction factor $\mathrm{d}(pK_a)/\mathrm{d}T$ should be used to calculate the appropriate $pH_{\text{prep}}$ for each temperature experiment, so that $pH_{\text{target}}$ for each experiment is constant across the entire range of temperatures probed. $\mathrm{d}(pK_a)/\mathrm{d}T$ values for common biological buffers have been reported in (*16, 17*). All buffers are prepared using Orion ROSS PerpHecT Micro Glass Bodied Combination pH Electrode (Thermo Scientific Cat. No. 8220BNWP) paired with Orion Temperature MicroATC probe (Thermo Scientific Cat. No. 928007MD).

Example calculation

To make a $pH$ 6.5 folding buffer for a 75°C reaction at room temperature (21 °C), we calculate nominal $pH_{\text{prep}}$ as follows:

$$\begin{aligned} pH_{\text{prep}} &= pH_{\text{target}} - \left(T_{\text{target}} - T_{\text{prep}}\right) \times \frac{d(pK_a)}{dT} \\ &= 6.5 - (54°C) \times (-0.017°C^{-1}) \\ &= 7.418 \end{aligned}$$

Final $pH_{\text{prep}}$ for fourU thermometer folding buffer

| **Target Temp. (°C)** | **Room Temp. (°C)** | **ΔT (°C)** | **Final $pH_{\text{prep}}$** |
|---|---|---|---|
| 10 | 20.5 | -10.5 | 6.32 |
| 15 | 19.6 | -4.6 | 6.42 |
| 18 | 19.4 | -1.4 | 6.48 |
| 20 | 19.2 | 0.8 | 6.51 |
| 25 | 19.8 | 5.2 | 6.59 |
| 30 | 19.3 | 10.7 | 6.68 |
| 33 | 19 | 14 | 6.74 |
| 37 | 19.6 | 17.4 | 6.8 |
| 40 | 19.2 | 20.8 | 6.85 |
| 42 | 20.2 | 21.8 | 6.88 |
| 45 | 21 | 24 | 6.92 |

| | | | |
|---|---|---|---|
| 48 | 18.8 | 29.2 | 6.98 |
| 65 | 19.1 | 45.9 | 7.29 |
| 70 | 19.7 | 50.3 | 7.36 |
| 75 | 18.7 | 56.3 | 7.45 |
| 80 | 18.9 | 61.1 | 7.54 |

Final $pH_{prep}$ for HIV-1 TAR folding buffer

| **Target Temp. (°C)** | **Room Temp. (°C)** | **ΔT (°C)** | **Final $pH_{prep}$** |
|---|---|---|---|
| 25 | 22.0 | 5.5 | 6.44 |
| 70 | 22.4 | 5.1 | 7.21 |
| 75 | 23.3 | 5.6 | 7.28 |
| 80 | 22.3 | 6.1 | 7.38 |

Final $pH_{prep}$ for P4-P6 folding buffer

| **Target Temp. (°C)** | **Room Temp. (°C)** | **ΔT (°C)** | **Final $pH_{prep}$** |
|---|---|---|---|
| 23 | 20.1 | 2.9 | 8.05 |

**Table S1**. Full sequences for RNA constructs used in this study. `Purple CAPITAL`: target RNA, **bold/underlined**: native target start/end, `lowercase`: primer binding site, `red-highlighted`: mutation sites.

| **Construct** | **Variant** | **RNA sequence** | **Source** | **Target start** | **Target end** | **Flank seq. design** |
|---|---|---|---|---|---|---|
| *Salmonella* fourU | WT | gguguaagggugaaguguaAGGUUGAACUUUUGAAUAGUG AUUCAGGAGGUUAAUGGAAguaaagguaaugaaggugaag | (*18*) | 2G | 36U | NUPACK |
| *Salmonella* fourU | A8C | gguguaagggugaaguguaAGGUUGACCUUUUGAAUAGUG AUUCAGGAGGUUAAUGGAAguaaagguaaugaaggugaag | (*18*) | 2G | 36U | NUPACK |
| HIV-1 TAR | WT | ggcaccucauaacauaacUAAGGCAGAUCUGAGCCUGGGA GCUCUCUGCCAAUCCacuaaccucacucacaauc | (*4*) | 17G | 45C | NUPACK |
| HIV-1 TAR | UUCG | ggcaccucauaacauaacUAAGGCAGAUCUGAGCUUCGGC UCUCUGCCAAUCCacuaaccucacucacaauc | (*7*) | 17G | 45C | NUPACK |
| HIV-1 TAR | A35G | ggcaccucauaacauaacUAAGGCAGAUCUGAGCCUGGGG GCUCUCUGCCAAUCCacuaaccucacucacaauc | (*4*) | 17G | 45C | NUPACK |
| HIV-1 TAR | C30U | ggcaccucauaacauaacUAAGGCAGAUCUGAGCUUGGGA GCUCUCUGCCAAUCCacuaaccucacucacaauc | (*4*) | 17G | 45C | NUPACK |
| HIV-1 TAR | UUCG-ES2 | ggcaccucauaacauaacUAAGGCAGAUCUGAGCCUUCGG GAGCUCUCUGCCAAUCCacuaaccucacucacaauc | (*6*) | 17G | 45C | NUPACK |
| *Tetrahymena* P4-P6 | WT | ggauaauccaaaacaacaaccAAAGGAAUUGCGGGAAAGG GGUCAACAGCCGUUCAGUACCAAGUCUCAGGGGAAACUUU GAGAUGGCCUUGCAAAGGGUAUGGUAAUAAGCUGACGGAC AUGGUCCUAACCACGCAGCCAAGUCCUAAGUCAACAGAUC UUCUGUUGAUAUGGAUGCAGUUCAACCAAAUCAccaaacc aaagaaacaacc | (*19, 20*) | 104A | 261A | Custom |

**Table S2.** gBlock constructs used in this study. Gray: padded 5' upstream sequence, green: T7 promoter, black: transcribed RNA. Note the transcribed RNA contains two guanine residues from T7 transcription.

| **Name** | **Sequence** | **Primers used** |
|---|---|---|
| fourU_WT_gblock | GTAAAACGACGGCCAGTGAATTCGAGCTCGGTACCTAATA<br>CGACTCACTATAGGTGTAAGGGTGAAGTGTAAGGTTGAAC<br>TTTTGAATAGTGATTCAGGAGGTTAATGGAAGTAAAGGTA<br>ATGAAGGTGAAG | DNA_template_F, fourU_DNA_R |
| fourU_A8C_gblock | GTAAAACGACGGCCAGTGAATTCGAGCTCGGTACCTAATA<br>CGACTCACTATAGGTGTAAGGGTGAAGTGTAAGGTTGACC<br>TTTTGAATAGTGATTCAGGAGGTTAATGGAAGTAAAGGTA<br>ATGAAGGTGAAG | DNA_template_F, fourU_DNA_R |
| HIV_WT_gblock | GTAAAACGACGGCCAGTGAATTCGAGCTCGGTACCTAATA<br>CGACTCACTATAGGCACCTCATAACATAACTAAGGCAGAT<br>CTGAGCCTGGGAGCTCTCTGCCAATCCACTAACCTCACTC<br>ACAATC | DNA_template_F, HIV_DNA_R |
| HIV_GS_UUCG_gblock | AGTAAAACGACGGCCAGTGAATTCGAGCTCGGTACCTAAT<br>ACGACTCACTATAGGCACCTCATAACATAACTAAGGCAGA<br>TCTGAGCTTCGGCTCTCTGCCAATCCACTAACCTCACTCA<br>CAATC | DNA_template_F, HIV_DNA_R |
| HIV_ES1_C30U_gblock | GTAAAACGACGGCCAGTGAATTCGAGCTCGGTACCTAATA<br>CGACTCACTATAGGCACCTCATAACATAACTAAGGCAGAT<br>CTGAGCTTGGGAGCTCTCTGCCAATCCACTAACCTCACTC<br>ACAATC | DNA_template_F, HIV_DNA_R |
| HIV_ES1_A35G_gblock | GTAAAACGACGGCCAGTGAATTCGAGCTCGGTACCTAATA<br>CGACTCACTATAGGCACCTCATAACATAACTAAGGCAGAT<br>CTGAGCCTGGGGGCTCTCTGCCAATCCACTAACCTCACTC<br>ACAATC | DNA_template_F, HIV_DNA_R |
| HIV_ES2_UUCG_gblock | GTAAAACGACGGCCAGTGAATTCGAGCTCGGTACCTAATA<br>CGACTCACTATAGGCACCTCATAACATAACTAAGGCAGAT<br>CTGAGCCTTCGGGAGCTCTCTGCCAATCCACTAACCTCAC<br>TCACAATC | DNA_template_F, HIV_DNA_R |
| P4P6_WT | GTAAAACGACGGCCAGTGAATTCGAGCTCGGTACCTAATA<br>CGACTCACTATAGGATAATCCAAAACAACAACCAAAGGAA<br>TTGCGGGAAAGGGGTCAACAGCCGTTCAGTACCAAGTCTC<br>AGGGGAAACTTTGAGATGGCCTTGCAAAGGGTATGGTAAT<br>AAGCTGACGGACATGGTCCTAACCACGCAGCCAAGTCCTA<br>AGTCAACAGATCTTCTGTTGATATGGATGCAGTTCAACCA<br>AATCACCAAACCAAAGAAACAACC | DNA_template_F, P4P6_DNA_R |

**Table S3**. Time-course chemical probing experiment overview with temperature and replicates for each construct.

| Construct | Variant | Temperature in °C (replicates) |
|---|---|---|
| *Salmonella* fourU | WT | 10(2), 15(2), 18(2), 20(2), 25(3), 30(4), 33(2), 37(5), 45(2), 52(2), 60(2), 65(2), 70(4), 75(4), 80(4) |
| *Salmonella* fourU | A8C | 15(2), 20(2), 25(4), 30(4), 37(3), 40(2), 42(2), 45(4), 48(2), 52(2), 60(2), 70(2), 75(2), 80(2) |
| HIV-1 TAR | WT | 25(2), 70(2), 75(2), 80(2) |
| HIV-1 TAR | A35G | 25(2), 70(2), 75(2), 80(2) |
| HIV-1 TAR | C30U | 25(2), 70(2), 75(2), 80(2) |
| HIV-1 TAR | UUCG | 25(2), 70(2), 75(2), 80(2) |
| HIV-1 TAR | UUCG-ES2 | 25(2), 70(2), 75(2), 80(2) |
| *Tetrahymena* P4-P6 | WT | 23-NoMg (1+2 endpoint replicates), 23-5mMMg(1) |

**Table S4**. Primers used for DNA template amplification, reverse transcription (RT), and library preparation. 'm' indicates 2'-O-methyl modification, which is included to minimize 3'-heterogeneity during T7 transcription (*21*). 'PCR1_F' and 'PCR1_R' are forward and reverse primers, respectively, for first round cDNA PCR. 'PCR2_i5_F' and 'PCR_i7_R' are Illumina indexing primers used in second round amplification. Asterisk (*) represents phosphorothioated bond to protect the primer from exonuclease degradation, retained from a prior PCR cleanup protocol not used in this study.

| Name | Step | Sequence | Purification |
|---|---|---|---|
| DNA_template_F | DNA template | GTAAAACGACGGCCAGTGAATTCG | Standard desalting |
| fourU_DNA_R | DNA template | mCmUTCACCTTCATTACCTTTACTTCCATTAA CC | Standard desalting |
| fourU_RT_primer | RT | CTTCACCTTCATTACCTTTA | Ion Exchange HPLC |
| fourU_PCR1_F | Library Prep | CTTTCCCTACACGACGCTCTTCCGATCTRRRYCTTCACCTTCATTACCTTTA*C *T*T*C | PAGE |
| fourU_PCR1_R | Library Prep | TGAACAGCGACTAGGCTCTTCAGGTGTAAGGGTGAAGTGTA | PAGE |
| HIV_DNA_R | DNA template | mGmATTGTGAGTGAGGTTAGTGGATTGGC | Standard Desalting |
| HIV_RT_primer | RT | GATTGTGAGTGAGGTTAGT | PAGE |
| HIV _PCR1_F | Library Prep | CTTTCCCTACACGACGCTCTTCCGATCTRRRYGATTGTGAGTGAGGTTAGT*G* G | PAGE |
| HIV_PCR1_R | Library Prep | TGAACAGCGACTAGGCTCTTCAGGCACCTCATAACATAAC | PAGE |
| P4P6_DNA_R | DNA template | mGmGTTGTTTCTTTGGTTTGGTTTT | Standard desalting |
| P4P6_RT_primer | RT | GGTTGTTTCTTTGGTTTGG | PAGE |
| P4P6_PCR1_F | Library Prep | CTTTCCCTACACGACGCTCTTCCGATCTNNNNGGTTGTTTCTTTGGTTTGGTTT | PAGE |
| P4P6_PCR1_R | Library Prep | TGAACAGCGACTAGGCTCTTCAGGATAATCCAAAACAACAACC | PAGE |
| PCR2_i7_R | Library Prep | CAAGCAGAAGACGGCATACGAGAT[8nt_i7]GTGACTGGAGTTCAGACGTGTG CTCTTCCGATCTTGAACAGCGACTAGGCTCTTCA | Ultramer |
| PCR2_i5_F | Library Prep | AATGATACGGCGACCACCGAGATCTACAC[8nt_i5]ACACTCTTTCCCTACAC GACGCTCTTCCGATCT | Ultramer |

**Table S5**. NMR kinetic measurement scan parameters. A600: Bruker Avance III 600 MHz system, HFCN600: Bruker Neo 600 MHz system

| **Sample** | **Temperature in °C (replicates)** | **Number of scans** | **Time per read (mins)** | **Total reads (time)** | **Machine** |
|---|---|---|---|---|---|
| DMS | 25 (2) | 64 | 5 | 96 (8 hrs) | A600 |
| DMS | 30 (2) | 64 | 5 | 60 (5 hrs) | A600 |
| DMS | 37 (2) | 32 | 2.5 | 36 (90 mins) | A600 |
| DMS | 45 (2) | 16 | 1.5 | 30 (45 mins) | A600 |
| DMS | 52 (2) | 16 | 1.5 | 20 (30 mins) | A600 |
| DMS | 60 (2) | 16 | 1.5 | 10 (15 mins) | A600 |
| ATP + DMS | 20 (2) | 152 | 10 | 60 (10 hrs) | A600 |
| ATP + DMS | 37 (2) | 32 | 2.5 | 36 (90 mins) | A600 |
| ATP + DMS | 42 (2) | 22 | 2 | 24 (44 mins) | A600 |
| ATP + DMS | 48 (2) | 16 | 1.5 | 20 (30 mins) | A600 |
| CTP + DMS | 25 (2) | 152 | 10 | 60 (10 hrs) | A600 |
| CTP + DMS | 37 (2) | 32 | 2.5 | 36 (90 mins) | A600 |
| CTP + DMS | 42 (2) | 22 | 2 | 24 (44 mins) | A600 |
| CTP + DMS | 48 (2) | 16 | 1.5 | 20 (30 mins) | A600 |
| GTP + DMS | 25 (2) | 80 | 10 | 60 (10 hrs) | HFCN600 |
| GTP + DMS | 33 (2) | 36 | 5 | 24 (2 hrs) | HFCN600 |
| GTP + DMS | 42 (2) | 12 | 2 | 24 (48 mins) | HFCN600 |
| GTP + DMS | 48 (1) | 16 | 1.5 | 20 (30 mins) | A600 |

**Table S6**. Methylation-induced chemical shift changes in ATP, CTP, GTP, and UTP. Prime symbol (') represent upfield shift of the reporter site as a result of methylation.

| NTP | Replicate Name | Temp (°C) | DMS (ppm) | $C_a$' (ppm) | $C_a$ (ppm) | Identity | $C_b$' (ppm) | $C_b$ (ppm) | Identity |
|---|---|---|---|---|---|---|---|---|---|
| ATP | 20_1 | 20 | 3.841 | 5.968 | 5.875 | C1' | 8.153 | 7.970 | C8 |
| ATP | 37_1 | 37 | 4.036 | 6.152 | 6.085 | C1' | 8.288 | 8.226 | C8 |
| ATP | 37_2 | 37 | 4.037 | 6.151 | 6.078 | C1' | 8.289 | 8.233 | C8 |
| ATP | 42_1 | 42 | 4.094 | 6.204 | 6.137 | C1' | 8.352 | 8.284 | C8 |
| ATP | 42_2 | 42 | 4.092 | 6.200 | 6.131 | C1' | 8.347 | 8.280 | C8 |
| ATP | 48_1 | 48 | 4.157 | 6.267 | 6.187 | C1' | 8.413 | 8.332 | C8 |
| ATP | 48_1 | 48 | 4.158 | 6.266 | 6.197 | C1' | 8.419 | 8.317 | C8 |
| GTP | 25_1 | 25 | 3.903 | 6.045 | 5.689 | C1' | 8.130 | 7.869 | C8 |
| GTP | 33_1 | 33 | 3.996 | 5.952 | 5.785 | C1' | 8.204 | 7.959 | C8 |
| GTP | 42_1 | 42 | 4.097 | 6.038 | 5.860 | C1' | 8.251 | 8.065 | C8 |
| GTP | 48_1 | 48 | 4.161 | 6.115 | 5.918 | C1' | 8.325 | 8.064 | C8 |
| CTP | 25_1 | 25 | 3.896 | 6.112 | 5.925 | C5 | 7.937 | 7.777 | C6 |
| CTP | 37_1 | 37 | 4.034 | 6.204 | 6.073 | C5 | 8.016 | 7.918 | C6 |
| CTP | 37_2 | 37 | 4.037 | 6.208 | 6.071 | C5 | 8.021 | 7.932 | C6 |
| CTP | 42_1 | 42 | 4.089 | 6.265 | 6.131 | C5 | 8.122 | 7.869 | C6 |
| CTP | 42_2 | 42 | 4.092 | 6.291 | 6.117 | C5 | 8.095 | 7.902 | C6 |
| CTP | 48_1 | 48 | 4.157 | 6.328 | 6.205 | C5 | 8.133 | 8.009 | C6 |
| CTP | 48_1 | 48 | 4.161 | 6.324 | 6.193 | C5 | 8.177 | 7.929 | C6 |
| UTP | 20_1 | 20 | 3.849 | | | | 7.887 | 7.691 | C6 |
